# A deep lung cell atlas reveals cytokine-mediated lineage switching of a rare cell progenitor of the human airway epithelium

**DOI:** 10.1101/2023.11.28.569028

**Authors:** Avinash Waghray, Isha Monga, Brian Lin, Viral Shah, Michal Slyper, Bruno Giotti, Jiajie Xu, Julia Waldman, Danielle Dionne, Lan T. Nguyen, Wendy Lou, Peiwen Cai, Eric Park, Christoph Muus, Jiawei Sun, Manalee V Surve, Lujia Cha Cha Yang, Orit Rozenblatt-Rosen, Toni M Dolerey, Srinivas Vinod Saladi, Alexander M Tsankov, Aviv Regev, Jayaraj Rajagopal

**Author notes:** co-first authors (Avinash Waghray, Isha Monga, Brian Lin, Viral Shah, Michal Slyper, Bruno Giotti). co-senior authors; to whom correspondence should be addressed (A.T.), (A.R.), (J.R.).

## Abstract

The human airway contains specialized rare epithelial cells whose roles in respiratory disease are not well understood. Ionocytes express the Cystic Fibrosis Transmembrane Conductance Regulator (CFTR), while chemosensory tuft cells express asthma-associated alarmins. However, surprisingly, exceedingly few mature tuft cells have been identified in human lung cell atlases despite the ready identification of rare ionocytes and neuroendocrine cells. To identify human rare cell progenitors and define their lineage relationship to mature tuft cells, we generated a deep lung cell atlas containing 311,748 single cell RNA-Seq (scRNA-seq) profiles from discrete anatomic sites along the large and small airways and lung lobes of explanted donor lungs that could not be used for organ transplantation. Of 154,222 airway epithelial cells, we identified 687 ionocytes (0.45%) that are present in similar proportions in both large and small airways, suggesting that they may contribute to both large and small airways pathologies in CF. In stark contrast, we recovered only 3 mature tuft cells (0.002%). Instead, we identified rare bipotent progenitor cells that can give rise to both ionocytes and tuft cells, which we termed tuft-ionocyte progenitor cells (TIP cells). Remarkably, the cycling fraction of these TIP cells was comparable to that of basal stem cells. We used scRNA-seq and scATAC-seq to predict transcription factors that mark this novel rare cell progenitor population and define intermediate states during TIP cell lineage transitions en route to the differentiation of mature ionocytes and tuft cells. The default lineage of TIP cell descendants is skewed towards ionocytes, explaining the paucity of mature tuft cells in the human airway. However, Type 2 and Type 17 cytokines, associated with asthma and CF, diverted the lineage of TIP cell descendants *in vitro*, resulting in the differentiation of mature tuft cells at the expense of ionocytes. Consistent with this model of mature tuft cell differentiation, we identify mature tuft cells in a patient who died from an asthma flare. Overall, our findings suggest that the immune signaling pathways active in asthma and CF may skew the composition of disease-relevant rare cells and illustrate how deep atlases are required for identifying physiologically-relevant scarce cell populations.

## MAIN

Single cell profiling studies in human and mouse airways have revealed the existence and gene expression profiles of a triad of rare cells: ionocytes, tuft cells, and neuroendocrine (NE) cells^1–11^. Ionocytes express the Cystic Fibrosis Transmembrane Conductance Regulator (CFTR), the causal gene whose disruption leads to cystic fibrosis (CF) ^6,7^. Indeed, their ablation in a ferret model of CF has recently been shown to result in abnormalities of airway physiology remarkably reminiscent of CF, and their loss in human large airway epithelial cultures disrupts ion transport ^12–14^. In contrast, tuft cells and NE cells have roles in environmental sensing and initiating inflammatory responses ^7,15^. However, the precise functions of these recently described rare cells and the signaling cascades that regulate their differentiation are not well understood. In particular, despite the generally accepted role of tuft cells in type 2 mediated inflammation in the intestine and in allergic rhinitis, exceedingly few tuft cells have been found in normal human airways^5,16^. “Ectopic” tuft cells have been reported in rare lung diseases, such as desquamative interstitial pneumonitis^17^ and immotile cilia syndrome^18^, as well as in lungs collected post mortem from individuals with COVID-19^19^. Moreover, although tuft cells participate in murine Th2-mediated allergic asthma and intestinal parasitic infection^20–23^, mature tuft cells have surprisingly not been detected in human asthmatic airways^3^. Similarly, ionocytes have been reported to be very rare in human small airways, despite the prominent small airway pathology seen in CF^6,7,24^. This has led to the suggestion that ionocytes may not play a significant role in the small airways pathology characteristic of the early stages of CF^24^.

To systematically address this challenge, we constructed an anatomically-guided, deep single cell RNA-Seq (scRNA-seq) atlas of 311,748 cells from 8 donor lungs, including 71,263 cells sampled from 14 regions of a single donor lung. We computationally reconstructed cell states during the differentiation of rare epithelial cells from their progenitors and identified a highly replicating bipotent progenitor cell population predicted to give rise to either tuft cells or ionocytes, that we accordingly termed Tuft-Ionocyte Progenitor (TIP) cells. Thus, in addition to the classic airway basal stem cell that sits atop the entire airway epithelial hierarchy, the human airway epithelium contains a second highly replicative progenitor cell population further down the lineage tree that is dedicated to the production of rare epithelial cells and is also capable of lineage switching. We coupled an optimized *in vitro* culture system to a new sorting methodology to enrich and profile rare cells that had been subjected to cytokine-mediated rare cell lineage skewing. We demonstrate that Type 2 and Type 17 cytokines, associated with asthma and CF, divert TIP cell lineage to result in the differentiation of mature tuft cells at the expense of ionocytes.

### Deep cell atlas of the human lung

To build a deep human lung cell atlas, we profiled 311,748 cells from 8 normal explanted human lungs, from individuals of both genders (5 females, 3 males), a broad age range (3 months to 66 years old) and different ancestries (6 white, one African American, and one Hispanic/Latino), representing the largest single dataset of normal human lung cells profiled to date (**Fig. 1a,b**, **Extended Data Fig. 1-3**, **Extended Table 1**). From one individual (HU37), we profiled 71,263 single cells across 13 anatomic sites along the respiratory tree, including 6 proximal airway sites and 7 distal lung lobe sites (**Fig. 1a,b**, **Extended Table 1**). We complemented this dataset with cells obtained from another 7 donors to account for inter-individual variability (**Extended Data Fig. 1-3**, **Extended Data Table 1**). The cells partitioned into 17 epithelial (**Fig. 1c,d** and **Extended Data Fig. 1**), 9 stromal/mesenchymal (**Extended Data Fig. 2a-d**), 5 endothelial (**Extended Data Fig. 2e-h**), and 32 immune (**Extended Data Fig. 3**) subsets, which were independently annotated and then matched to the Human Lung Cell Atlas (HLCA) ontology^16^.

**Figure 1.**
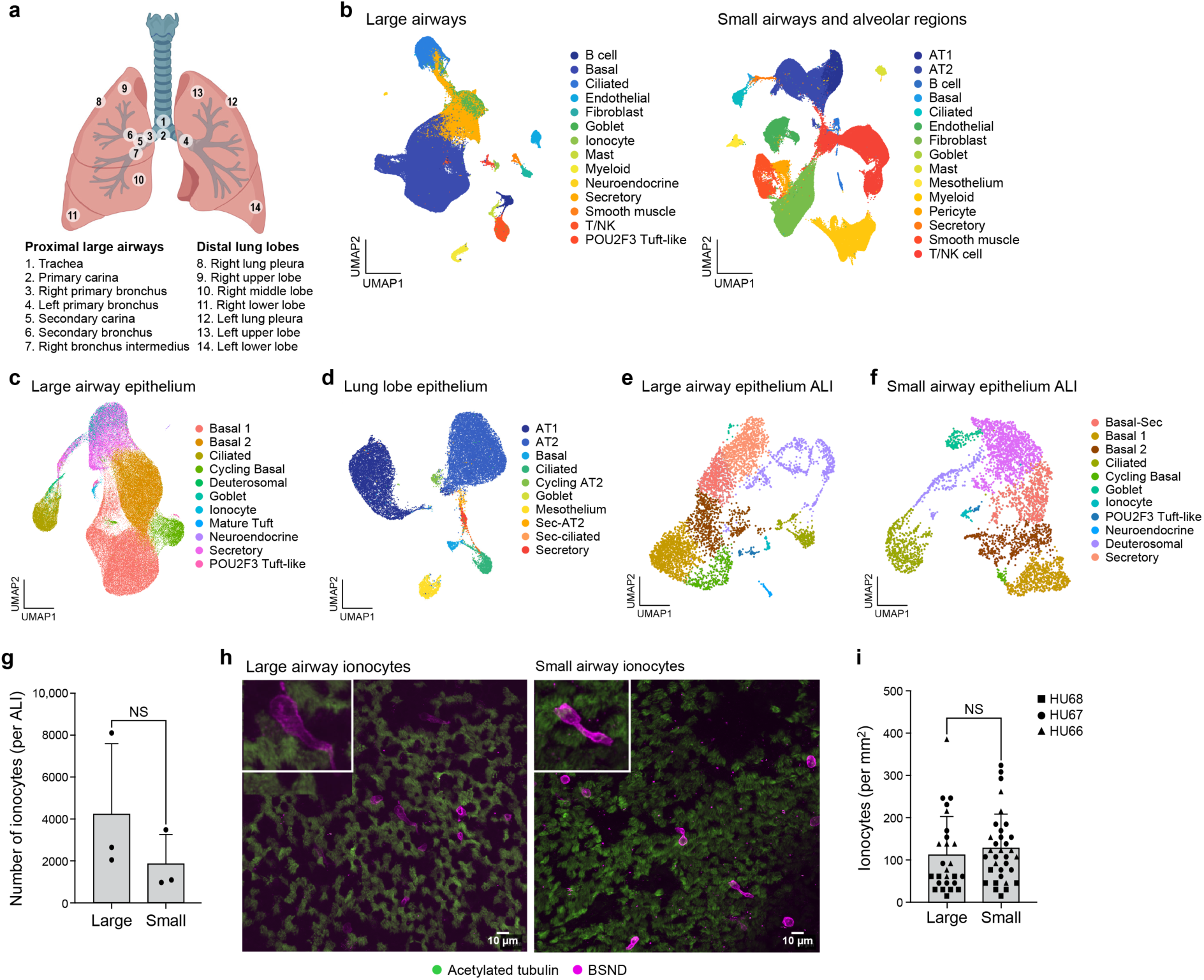
Deep cell atlas of the human lung reveals CFTR^+^ ionocytes in both human proximal and distal airways and proximal and distal ALI cultures. **a.** Regional sampling for a deep lung cell atlas. Numbered circles: sampled locations. **b.** Lung cell atlas. Uniform manifold approximation and projection (UMAP) embedding of cell profiles (dots) from the large airways (left) and lung lobe regions (right) colored by cell type annotation. **c-f.** Epithelial lung and ALI cell profiles. UMAP embeddings of epithelial cell profiles from the proximal airway (**c**), distal lung lobe (**d**), and ALI cultures generated from large airway basal cells isolated from primary bronchus (LAE ALI) (**e**) or from small airway basal cells isolated from primary bronchus (SAE ALI) (**f**). **g-i**. CFTR^+^ ionocytes in the human proximal and distal airways and human proximal and distal ALI cultures. **g**. Number of BSND^+^ mature ionocytes per ALI (y axis) in LAE ALI and SAE ALI cultures (x axis). n = 3, Two tailed unpaired T test. **h**. Whole mount images of dissected large (left) and small (right) airways stained for BSND (magenta) and acetylated Tubulin (green). Insets: BSND^+^ ionocytes (magenta). **i**. Number of BSND^+^ mature ionocytes per mm^2^ (y axis) in micro dissected large airways and small airways (x axis). n=3 on three normal human lungs (Hu66, Hu67 and Hu68). ANOVA (Sidak’s multiple comparisons).

**Figure 2.**
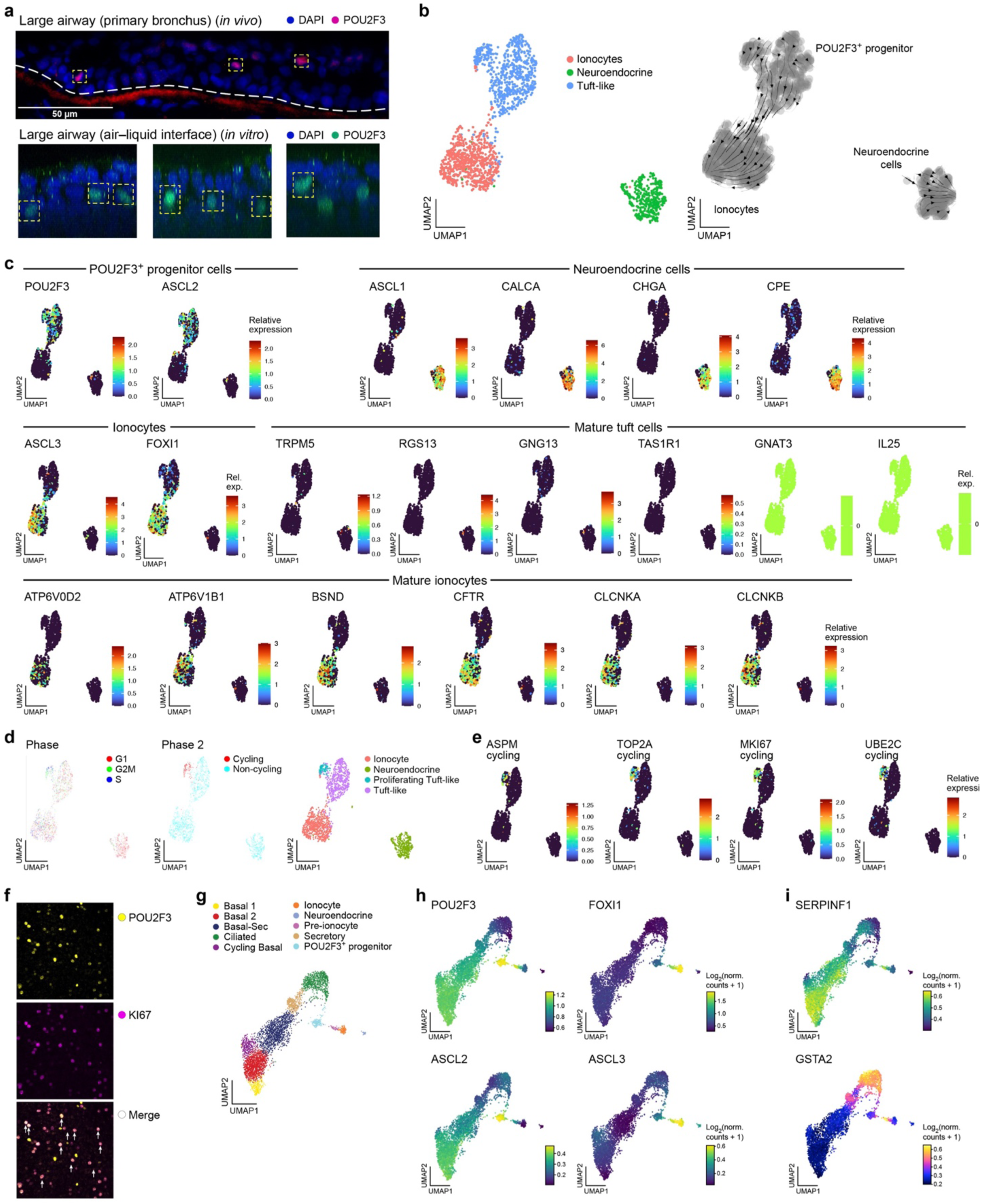
Expression and chromatin profiles define a replicative rare cell progenitor and a pre-ionocyte. **a.** Human large airways and human LAE ALI cultures both contain POU2F3^+^ cells. Antibody staining of a section of right primary bronchus with POU2F3 (red) and DAPI (blue) and LAE ALI with POU2F3 (green) and DAPI (blue). **b,c.** POU2F3^+^ cells are predicted to be progenitors of mature ionocytes. UMAP embedding of scRNA-seq profiles (dots) of rare cells, colored by annotation (b, left), labeled with RNA velocity analysis (b, right), or colored by expression of markers for different rare cell subsets (c) **d-f.** POU2F3^+^ cells have a sizable fraction of replicating cells. **d-e.** UMAP embedding of scRNA-seq profiles (dots) of rare cells, colored by cell cycle phase annotation (d, left), cycling gene signature based annotation (d, middle), POU2F3+ tuft-like cells are comprised of a replicating and a non-replicating fraction (d, right) as gauged by the expression of specific cell cycle genes (e) **f.** POU2F3^+^ (yellow) cells in LAE ALI cultures co-stained with MKI67 (purple). **g-i.** Distinct chromatin state marks POU2F3^+^ progenitors. UMAP embedding of large airways epithelial cell scATAC-seq profiles (dots) colored by cell type annotation (g) or accessibility at select gene loci (h,i).

**Figure 3.**
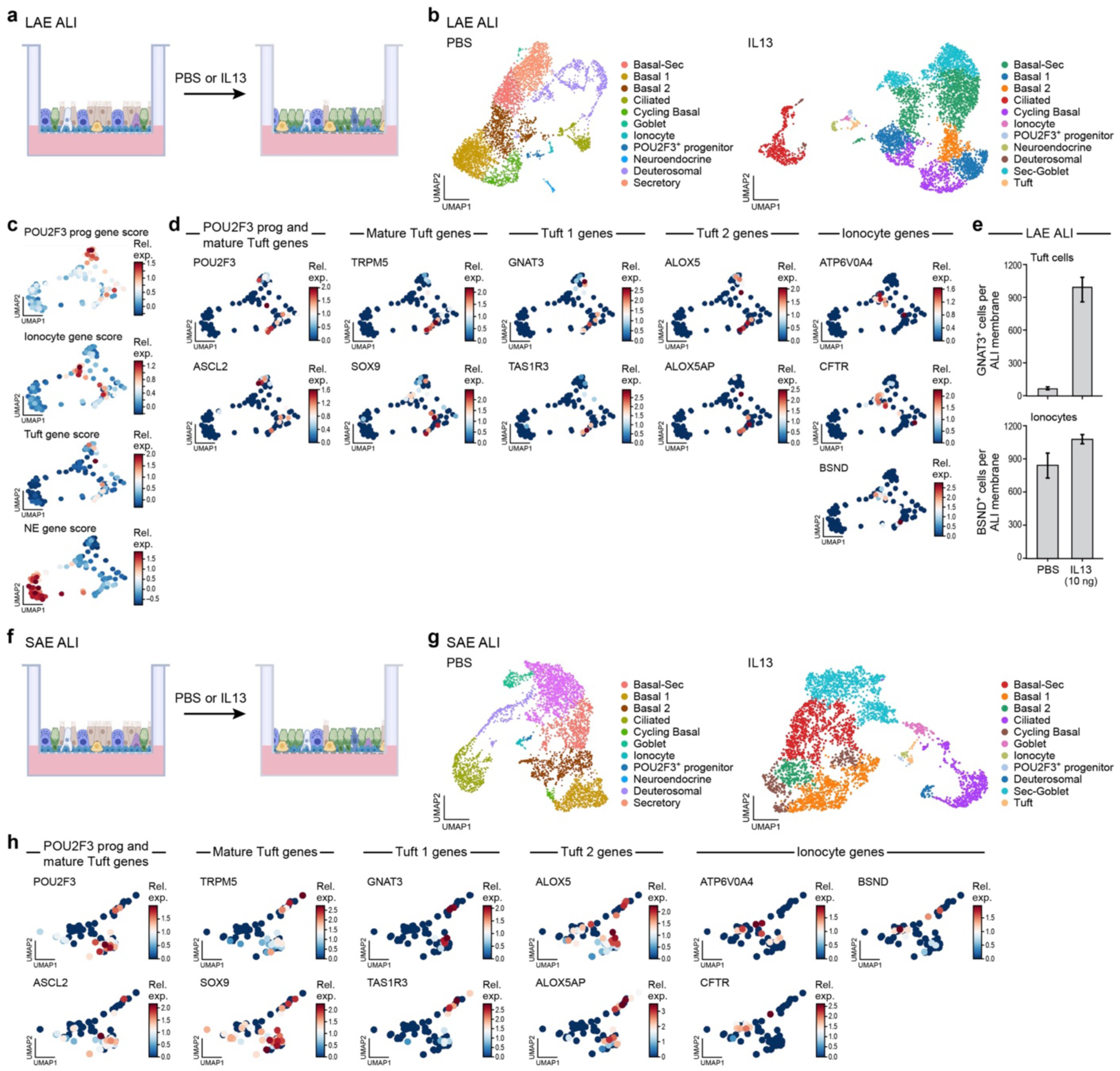
Type 2 pathway cytokines induce mature tuft cell differentiation. **a.** Overview of LAE ALI experimental system. Mature LAE ALIs are treated for 96 hours with PBS (control) or IL13 (20 ng/ml). **b.** IL-13 treatment shifts LAE ALI cell composition. UMAP embedding of scRNA-seq profiles (dots) from control (left; as in Fig. 1e) and IL-13 treated (right) LAE ALIs, colored by cell type annotation. **c-e.** Mature tuft cells are induced in IL-13 treated LAE ALIs. Zoom of portion of the UMAP embedding in IL-13 treated LAE ALIs (from **b**, right) colored by scores for rare cell marker gene signatures (**Extended Data Table 2**) (c) or by expression of rare cell marker genes (d). **e.** Number of antibody-stained cells (y axis) for GNAT3 expressing tuft cells (n=2 (Hu19, Hu67)) and BSND expressing ionocytes n=3 (Hu19, Hu62, Hu67) in LAE ALI treated with PBS or IL13 (10ng/ml) (x axis). (The same PBS control as in Fig. 5b and **Extended Data Fig. 9d**; all experimental treatments were done in parallel; **Methods**). **f.** Overview of SAE ALI experimental system. Mature SAE ALIs are treated for 96 hours with PBS (control) or IL13 (20 ng/ml). **g.** IL-13 treatment shifts SAE ALI cell composition. UMAP embedding of scRNA-seq profiles (dots) from control (left; as in Fig. 1f) and IL-13 treated (right) SAE ALIs, colored by cell type annotation. **h.** Mature tuft cells are induced in IL-13 treated SAE ALIs. Zoom in of a portion of the UMAP embedding in IL-13 treated SAE ALIs (from **g**, right) colored by expression of rare cell marker genes.

In the proximal airway we identified all of the expected common epithelial cell types (basal, secretory, goblet, deutrosomal^10^, and ciliated cells), along with cells from the glands (**Extended Data Fig. 1a**), and the three previously identified rare cell populations: neuroendocrine (NE) cells, ionocytes, and POU2F3^+^ cells (**Fig. 1b**) . Previously, these rare POU2F3^+^ cells had been identified as “tuft-like cells”^1,16^, in part because POU2F3 is required for tuft cell differentiation and maintenance^20,25^. In the distal region of the lung, we again found all the expected cell types, including alveolar type 2 (AT2) cells, alveolar type 1 (AT1) cells, and mesothelial cells, as well as small airway epithelial cell types, including basal, secretory, goblet, and ciliated cells. Additionally, in alveolar regions we detected the previously described rare secretory cell subsets: secretory-ciliated (SecCil) and secretory-AT2 (SecAT2) cells^26,27^ (**Fig. 1d** and **Extended Data Fig. 1b**). In contrast, despite the anatomical sampling and depth of our cell atlas, we did not detect rare small airway epithelial cell types, due to the abundance of alveolar type 1 and type 2 cells in samples of lung lobe.

### Mature tuft cells are exceedingly rare in normal human lungs and *in vitro* models

In order to determine the abundance of mature tuft cells in human lungs, we first re-clustered the 885 annotated rare epithelial cells in the core HLCA dataset which is composed of harmonized data from 11 independent atlases ^28^ (**Extended Data Fig. 4a-c**). We then developed a scoring approach to robustly discriminate NE cells, ionocytes, POU2F3^+^ “tuft-like” cells, and mature tuft cells, using markers from the dataset^5^ that identified the greatest number of mature tuft cells (10 cells) in a single study (**Methods, Extended Data Table 2**). Using this approach, we confirmed that the HLCA dataset contains 561 ionocytes (0.096% of all cells; 0.199% of all epithelial cells) and 159 NE cells (0.027% of all cells; 0.056% of all epithelial cells) (**Extended Data Fig. 4d**, **Extended Data Table 3**). However, the 165 cells that were annotated as tuft cells in HLCA, were, in fact, composed of two subsets: A major subset of 152 POU2F3^+^ “tuft-like” cells (0.026% of all cells; 0.054% of all epithelial cells), and a very minor subset of 13 mature tuft cells (0.002% of all cells; 0.005% of all epithelial cells, **Extended Data Fig. 4d**, **Extended Data Table 3**).

**Figure 4.**
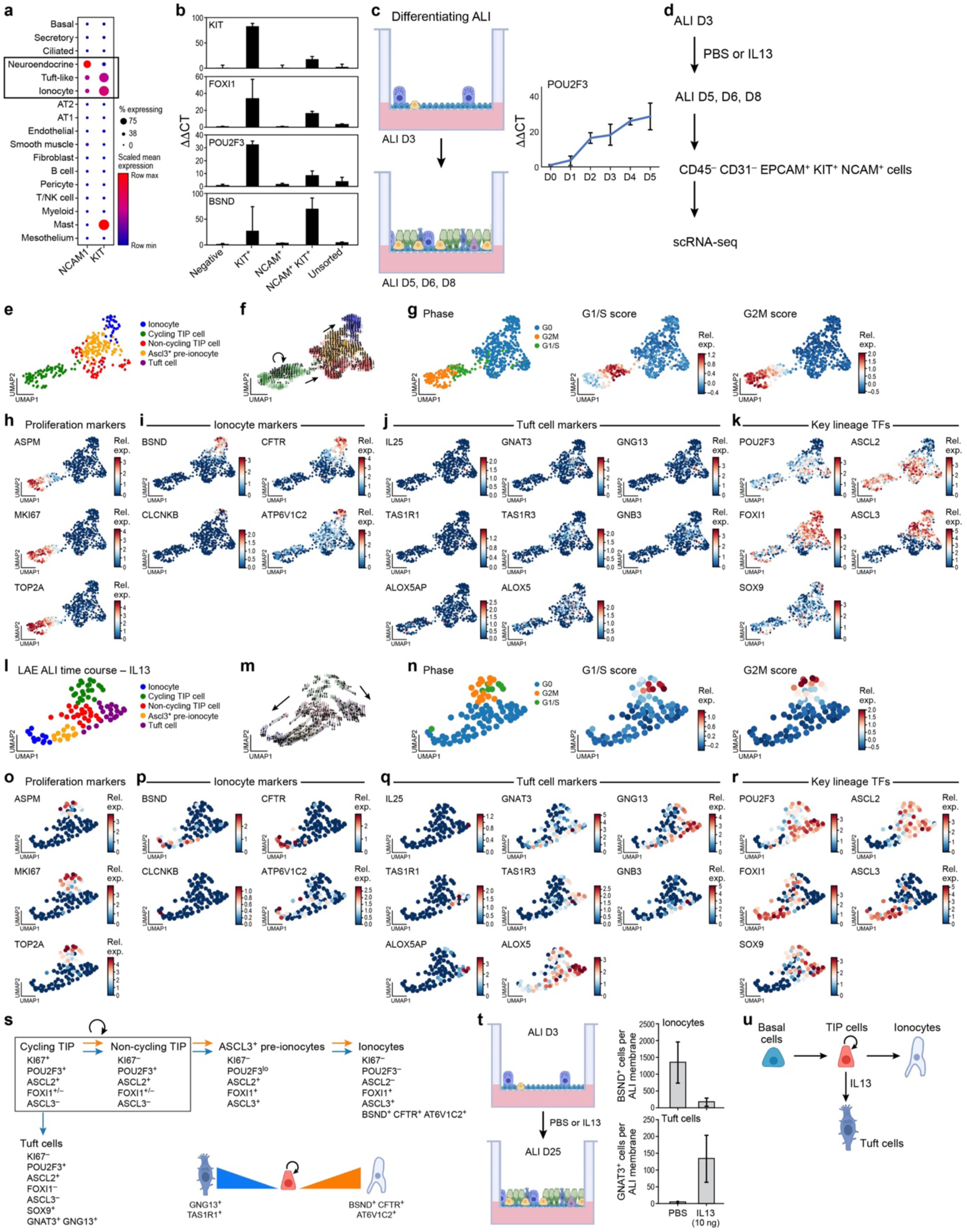
Type 2 pathway cytokines induce a lineage switch in POU2F3^+^ bipotent Tuft-Ionocyte Progenitor (TIP) cells. **a,b.** Sorting strategy to enrich rare cells. **a**. Mean expression (dot color, relative expression) and proportion of cells (dot size) expressing genes encoding the cell surface proteins NCAM1 and KIT in human *in vivo* scRNA-seq data of Fig. 1b. **b.** Expression level (∂∂CT, qPCR, y axis) of key rare cell marker genes (marked on top left) in sorted cell populations from human ALI cultures (x axis, labeled by sorting marker). n=3 technical replicates. **c,d.** Experimental strategy. **c.** Left: Model of differentiating ALI. Right: POU2F3 mRNA expression (∂∂CT, qPCR, y axis) at different time points (x axis) during ALI differentiation. n=3 technical replicates. **d.** Experimental time course. From top: starting at ALI D3 (the time point at which TIP cells are first present), cultures were treated with IL13 (10ng/ml) or PBS control for 5 days and then CD45^-^ CD31^-^ EPCAM^+^ NCAM1^+^, CD45^-^ CD31^-^ EPCAM^+^ KIT^+^ and CD45^-^ CD31^-^ EPCAM^+^ KIT^+^ NCAM1^+^ rare cells were collected, pooled, and profiled using scRNA-seq. **e-k.** TIP cells give rise to ionocytes via defined transitional states in control LAE ALI cultures. UMAP embedding of scRNA-seq profiles (dots) from PBS-treated (control) LAE ALI cultures colored by cell type annotation (e), overlaid RNA velocity vectors (f), cell cycle phase annotation (g, left), G1/S (g, middle) and G2/M (g, right) gene signature scores, and gene expression of markers of cell proliferation (h), ionocytes (i), tuft cells (j), and key lineage TFs (k). **l-r.** TIP cells give rise to both ionocytes and mature tuft cells following IL-13 treatment. UMAP embedding of scRNA-seq profiles (dots) from IL-13-treated LAE ALI cultures colored by cell type annotation (l), overlaid RNA velocity vectors (m), cell cycle phase annotation (n, left), G1/S (n, middle) and G2/M (n, right) gene signature scores, and gene expression of markers of cell proliferation (o), ionocytes (p), tuft cells (q), and key lineage TFs (r).**s.** Lineage model schematizing the expression of key TFs during rare cell differentiation. Orange arrows indicate PBS and blue arrows indicate IL13-induced differentiation**. t.** Left: Experimental setup of differentiating ALI treated continuously with IL-13 for 25 days. Right: Number of BSND^+^ cells (ionocytes, y axis, n=3 (Hu19, Hu60, Hu67)) and GNAT3^+^ cells (tuft cells, y axis, n=2 (Hu60, Hu67)) in PBS and IL-13 conditions (x axis). **u.** Model of cytokine-mediated TIP cell lineage switching.

Applying the same scoring approach to our deep atlas revealed 687 ionocytes (0.22% of all cells; 0.45% of airway epithelial cells), 363 NE cells (0.12% of all cells; 0.24% of airway epithelial cells), 725 POU2F3^+^ “tuft-like” cells (0.23% of all cells; 0.47% of airway epithelial cells), but only 3 mature tuft cells (0.001% of all cells; 0.002% of airway epithelial cells). Thus, despite our very deep sampling of rare cells, mature tuft cells were exceedingly rare in the normal human lung^5^ and do not form a distinct cluster using standard unsupervised community detection (**Extended Data Fig. 4e**; **Extended Data Table 4**). In total, we added 1,778 rare cell profiles to the 885 profiles contained in HLCA, and refined the HLCA annotation, establishing that mature tuft cells are a remarkably scarce fraction (<0.6%) of an already scarce total rare epithelial cell pool.

Given the exceedingly low proportion of rare epithelial cells recovered from dissociated distal lungs, we next compared the relative proportions of rare cells in large and small airway epithelial air-liquid interface cultures (LAE and SAE ALI, respectively). In contrast to prior reports that SAE ALI cultures contain fewer ionocytes than LAE ALIs^24^, scRNA-seq of LAE *vs*. SAE ALI cultures detected comparable numbers of ionocytes on average (per sampled ALI, **Methods**) (**Fig. 1e-g**). Notably. mature tuft cells were not detected in either LAE or SAE ALI cultures.

We then hypothesized that those physiological functions that are normally ascribed to mature tuft cells, such as environmental sensing, may be assumed by other cell types in the steady state human airway epithelium. Supporting this hypothesis, genes related to bitter taste receptors (*TAS2R4*, *TAS2R43*) are expressed by human ciliated cells^29^, genes related to olfactory receptor-mediated functions are expressed in NE^30^ cells and genes for succinate sensing (*e.g*., succinate receptor (*SUCNR1*)) are expressed in glandular NE cells^31^. Indeed, NE cells from our deep cell atlas and human LAE cultures expressed the succinate receptor (*SUCNR1*), whereas POU2F3^+^ “tuft-like” cells did not (**Extended Data Fig. 5a,b**). We confirmed SUCNR1 receptor protein expression in NE (TUJI^+^) cells in human LAE ALI culture *in situ* (**Extended Data Fig. 5c**) and further established that NE cells functionally respond to succinate with a rise in intracellular calcium (**Extended Data Fig. 5d**).

**Figure 5.**
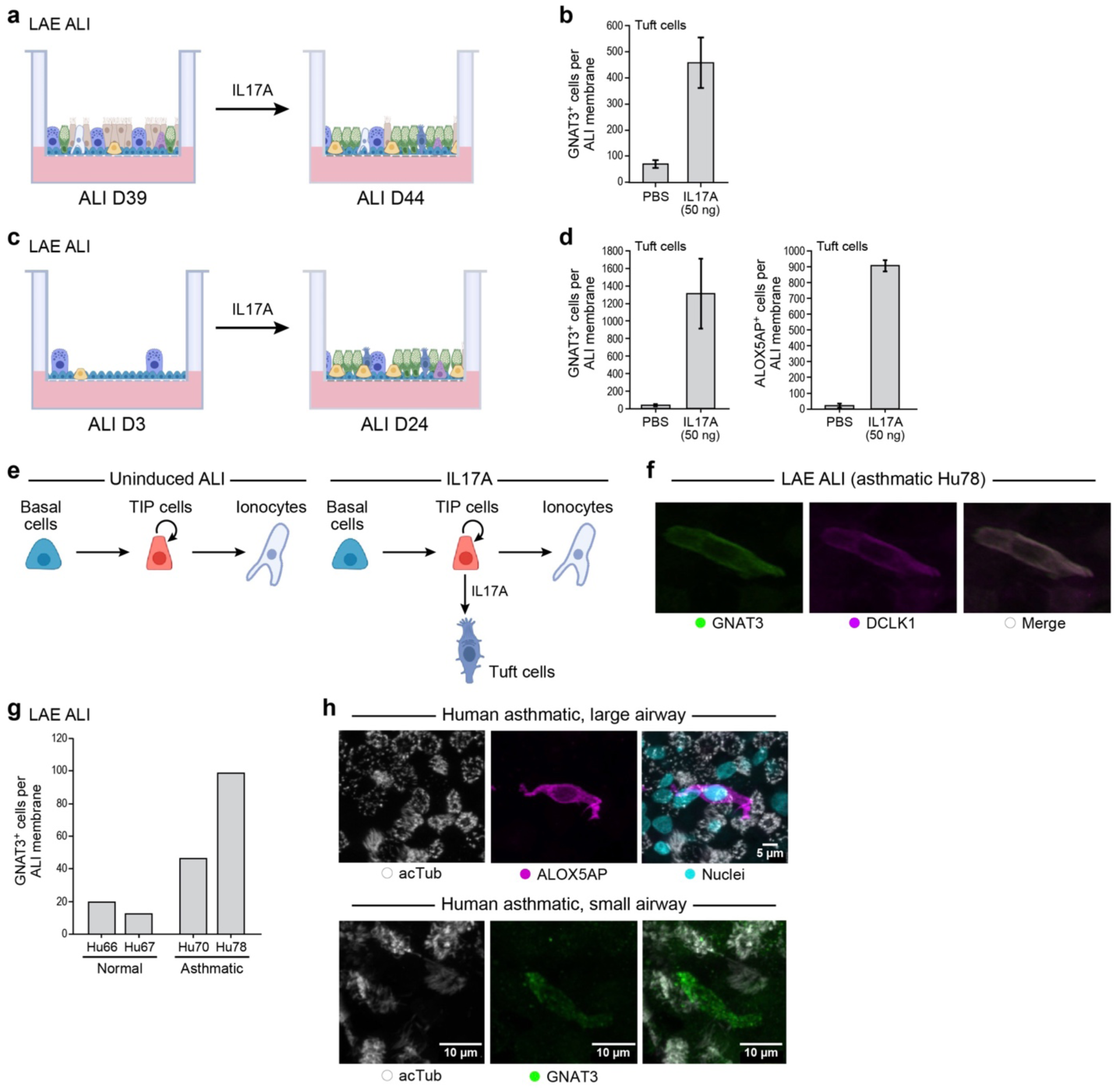
Type 17 signaling promotes tuft cell expansion at the cost of ionocyte differentiation and tuft cells are detected in asthmatic patient tissue sample. **a-d.** Tuft cell numbers are increased by IL-17 treatment. a,c. Overview of IL-17 treatment experiments in mature LAE ALI treated with PBS (control) or IL17-A (50ng/ml; from D39 to D44) (a) and differentiating ALI at D3 (the time point at which TIP cells are present) treated cells with IL17A (50ng/ml) or PBS from D3 to D24 (c). **b.** Number of GNAT3^+^ mature tuft cells (by immunohistochemistry, y axis, n=2 (Hu19, Hu67)) in PBS and IL17A (50ng/ul; from D39 to D44) treated mature LAE ALIs (x axis). (PBS control as in Fig. 3e and **Extended Data Fig. 9d**; the experimental treatments were done together.) **d.** Number of GNAT3^+^ mature tuft cells (by immunohistochemistry, y axis, n=2 (Hu19, Hu67)) and ALOX5AP^+^ mature tuft cells (by immunohistochemistry, y axis, n=2 (Hu19, Hu62)) of large airway ALIs treated with PBS or IL17A from D3 to D24. **e.** Model of Type 17 cytokine-induced TIP cell differentiation into mature tuft cells. **f-h.** Mature tuft cells in ALI and human tissue from asthmatic patients. **f.** GNAT3 (green) and DCLK1 (magenta) double positive mature tuft cells in LAE ALI generated from asthmatic patient Hu78. **g.** Number of GNAT3+ mature tuft cells in LAE ALIs derived from two asthmatic individuals (Hu70 and Hu78) and in patients with no history of lung diseases (Hu66, Hu67). **h.** Whole mount staining of dissected large airways and small airways from a patient with asthma exacerbation. The asthmatic patient’s large and small airway epithelia contain mature tuft cells (Hu78).

### CFTR-rich ionocytes are comparably distributed in large and small airways

Although CF is associated with small airways disease, prior reports claimed that small airways and SAE cultures contain few ionocytes^24,32^, leading to the suggestion that ionocytes do not play a role in CF-associated small airway pathology. In contrast, our LAE and SAE cultures contained a similar number of ionocytes (**Fig. 1g**). We further found that ionocyte abundance depended on the choice of culture media (**Extended Data Fig. 6a**), explaining the reported lack of *in vitro* ionocyte differentiation in prior studies^24^. In our cultures, both large airway and small airway ionocytes resemble their *in vivo* counterparts and comparably express ionocyte transcription factors (TFs) *FOXI1* and *ASCL3*^7^, *CFTR*, and the CF-associated ion transporters *ATP1A1*, *ATP6V0D2* and *ATP6V1C2* (**Extended Data Fig. 6b**). Finally, to directly quantify the numbers of sparsely distributed ionocytes *in situ* we performed whole mount staining of microdissected airways^24^ from three donors (**Extended Data Fig. 7**)^24^. We detected mature ionocytes expressing BSND protein in both large and small airways (less than 2 mm in diameter) (**Fig. 1h**) in similar numbers per epithelial surface area (**Fig. 1i**). Thus, small airway ionocytes expressing CF-related genes may yet play a role in CF-associated small airways disease^12^.

### Ionocytes arise from replicating POU2F3^+^ progenitor cells

POU2F3^+^ “tuft-like” cells were present in sections of primary bronchus as gauged by both scRNA-seq and *in situ* protein staining (**Fig. 2a**) and both lineage trajectory and RNA velocity analyzes ^33^ predicted that they would give rise to ionocytes (**Fig. 2b**). This prediction is consistent with prior observations that POU2F3 knockout results in the loss of ionocyte differentiation in large airway ALI cultures^1^. The POU2F3^+^ population expresses both ionocyte (*FOXI1*) and tuft cell (*POU2F3, ASCL2*) TFs (**Fig. 2c**). Furthermore, these cells do not express markers of mature tuft cells (*TRPM5, GNG13, RGS13, TAS1R1, GNAT3, IL25),* ionocytes (*BSND, CFTR, ATP6V0D2, ATP6V1B1, CLCNKA, CLCNKB*), nor of NE cells (*ASCL1, CALCA, CHGA, CPE*) (**Fig. 2c**). Taken together, these findings suggest that POU2F3^+^ cells, previously annotated as “tuft-like” cells ^1^, act as ionocyte progenitors during the course of normal homeostasis. RNA velocity analysis of cells profiled from SAE *vs*. LAE cultures suggest that both small and large airway ionocytes arise from POU2F3^+^ progenitor cells (**Extended Data Fig. 6c**).

Notably, a subset of human airway POU2F3^+^ progenitor cells (18.34%) were cycling *in vivo*, expressing canonical markers of cell proliferation (*KI67, ASPM, TOP2A, UBE2C*) and a cell cycle signature (**Fig. 2d,e**). Thus, the cycling fraction of POU2F3^+^ progenitor cells is greater than the cycling fraction of canonical basal stem cells (11%) in human large airways (**Extended data Table 1 and Table 7**). We confirmed the presence of proliferative POU2F3^+^ progenitor cells in ALI cultures (**Fig. 2f)**, and demonstrate that they also express *ASCL2,* mimicking their *in vivo* counterparts (**Extended Data Fig. 6d**). Interestingly, a fraction of POU2F3^+^ cells start to express the ionocyte TF *FOXI1* (**Fig. 2c**, **Extended Data Fig. 6d**). However, they do not express the full complement of TFs, such as *ASLC3*, present in mature ionocytes (**Fig. 2c**, **Extended Data Fig. 6d**).

### Chromatin accessibility profiles distinguish a pre-ionocyte state

To delineate the chromatin accessibility landscape underpinning tuft cell and ionocyte differentiation, we profiled 18,260 cells from Hu62 by single cell <u>A</u>ssay for <u>T</u>ransposase-<u>A</u>ccessible <u>C</u>hromatin with <u>S</u>equencing (scATAC-Seq). We identified 12 major large airway cell clusters (**Extended Data Fig. 8a,b**), including NE cells, ionocytes, and POU2F3^+^ progenitor cells (**Fig. 2g**), but not the exceedingly rare mature tuft cells (**Extended Data Table 5**), as well as 12 major cell clusters from the alveolar lung regions (**Extended Data Fig. 8c,d**). We then identified cell type specific TFs whose associated motifs were enriched in accessible regions of chromatin in each of the cell clusters (**Extended Data Fig. 8e,f**; ChromVar^34^ FDR≤0.01, mean differential accessibility ≥0.5). We further defined cell type specific open chromatin regions that can be used for cell type specific enhancer discovery and the construction of cell type specific reporters (**Extended Data Table 6**).

Interestingly, POU2F3^+^ cells partitioned into two clusters based on their scATAC-Seq profiles: a ‘POU2F3^+^ progenitor’ cluster and a ‘pre-ionocyte’ cluster (**Fig. 2g**). As expected, the POU2F3^+^ progenitors are characterized by high accessibility at the *POU2F3* and *ASCL2* loci (**Fig. 2h**, **Extended Data Fig. 8g**) suggesting a competence to differentiate into mature tuft cells. However, accessibility at the *POU2F3* and *ASCL2* loci gradually decreases as POU2F3^+^ progenitors transition into pre-ionocytes and then into ionocytes, alongside a concomitantly increasing accessibility at the ionocyte loci *FOXI1* and *ASCL3* (**Fig. 2h**, **Extended Data Fig. 8g**). This is consistent with commitment towards an ionocyte fate, prior to the opening of loci associated with mature ionocytes ^35^. Indeed, while some mature ionocyte gene loci (*CLCNKB, PTGER3*) remain largely closed in pre-ionocytes, other ionocyte loci (*BSND*) have already begun to open, and the *CFTR* locus is already decidedly open (**Extended Data Fig. 8h**). Interestingly, pre-ionocytes are also marked by unique open loci for genes such as *SERPINF1* and *GSTA2* that are not accessible in either POU2F3^+^ progenitor cells or mature ionocytes (**Fig. 2i**). In summary, we have identified committed transitional cells, pre-ionocytes, in which tuft cell TFs are being downregulated, while the mature ionocyte transcription factor program is being initiated.

### Type 2 cytokines IL-13 and IL-4 induce mature tuft cell differentiation

The coincident expression of *FOXI1* and *POU2F3* in the POU2F3^+^ progenitor cells that give rise to committed pre-ionocytes suggests the hypothesis that these same POU2F3^+^ progenitor cells are bipotent and that they can differentiate into either mature tuft cells or ionocytes depending on a particular signaling milieu. It is known that tuft cell-derived cysteinyl leukotrienes and the alarmin IL-25 are necessary for promoting Type 2 inflammation in murine models of asthma^23,36–38^. Reciprocally, the asthma-associated type 2 cytokine IL-13 induces tuft cell expansion in human and murine nasal epithelial cultures^39,40^. Remarkably, although tuft cells have been shown to be present in human allergic rhinosinusitis^40–42,40,43–45^, they have not been identified in asthma scRNA-seq datasets despite the combined analysis of 40,430 cells^3,46^.

To assess the role of Type 2 cytokines on human rare cell differentiation, we treated mature large airway epithelial air-liquid interface cultures (LAE ALI) with IL13 (20ng/μl) for 96 hours and then performed scRNA-seq (**Fig. 3a**). As expected^47^, IL13 treatment resulted in the differentiation of secretory cells into secretory-goblet cells (**Fig. 3b).** We continued to be able to identify POU2F3^+^ progenitors, ionocytes, and NE cells, but we also identified an additional new cluster of TRPM5^+^SOX9^+^POU2F3^+^ mature tuft cells (**Fig. 3c,d**), expressing markers of both murine tuft 1^7^ (chemosensory) airway epithelial cells (*GNAT3, TAS1R3*) and murine tuft 2^7^ (leukotriene) (*ALOX5, ALOX5AP*) cells (**Fig. 3d**). In further support of a model in which Type 2 cytokines induce bipotent POU2F3^+^ progenitor cells to differentiate into mature tuft cells, IL13 application results in the nascent expression of mature tuft cell genes (*TRPM5, GNAT3, ALOX5*) within the POU2F3^+^ progenitor cluster itself (**Fig. 3d)**. Immunohistochemistry confirmed that IL13 treatment resulted in a significant increase in GNAT3^+^ cells, while the number of ionocytes (BSND^+^) remained unchanged (**Fig. 3e**, **Extended data Fig. 9a-b**).

IL4, another Type 2 pathway cytokine, also induces the expression of mature tuft cell genes (*TRPM5, GNAT3*) in cultured LAE, as measured by qPCR of entire cultures, whereas treatment with the type 2 cytokine IL5 did not show a pronounced effect (**Extended Data Fig. 9c**). Immunohistochemistry confirmed that IL4 treatment, but not IL5 treatment, resulted in a significant increase in GNAT3^+^ tuft cells, while the number of ionocytes (BSND^+^) remained unchanged (**Extended Data Fig. 9d**). The lack of an effect of IL5 is consistent with the expression of the IL5 receptor (*IL5RA*) exclusively in ciliated and deutrosomal cells (**Extended Data Fig. 9e**). Treatment of small airway epithelial (SAE) ALI cultures with IL13 (20ng/μl) for 96 hours (**Fig. 3f**) also resulted in the differentiation of mature tuft cells (**Fig. 3g,h**).

### Type 2 cytokines induce a lineage switch in POU2F3^+^ bipotent tuft-ionocyte progenitor (TIP) cells

Although POU2F3^+^ progenitors express both tuft cell TFs, *POU2F3* and *ASCL2*, and were previously termed “tuft-like” cells^20,25,48^, we have shown that many POU2F3^+^ progenitors also express the ionocyte TF *FOXI1*^1^ and normally give rise to ionocytes (**Fig. 2b,c**). Indeed, chromatin accessibility at the *POU2F3* and *ASCL2* loci gradually decreases in pre-ionocytes concomitant with an increase in *FOXI1* accessibility (**Fig. 2h, Extended Data Fig. 8g**). Nonetheless, the coincident expression of tuft and ionocyte TFs suggests the hypothesis that POU2F3^+^ progenitors are bipotent and represent the source of IL13-induced mature tuft cells. To further examine the lineage trajectory leading to IL13-induced mature tuft cell differentiation, we developed an enrichment strategy to obtain a larger number of differentiating rare epithelial cells for subsequent computational analysis. In order to accomplish this, we used our *in vivo* scRNA-seq data to identify a combination of cell surface proteins that can be used for fluorescence activated cell sorting (FACS) of rare epithelial cells (**Fig. 4a**). Indeed, sorting for CD45^-^CD31^-^EPCAM^+^KIT^+^NCAM1^+^ cells effectively enriched for cells expressing *POU2F3* and *FOXI1* (**Fig. 4b**). Next, in order to enrich for the presence of actively differentiating progenitor cells, we opted to model lineage switching in a differentiating ALI culture system in lieu of the fully mature epithelial cultures we had previously used to demonstrate mature tuft cell differentiation (**Fig. 3a**). In this model system, *POU2F3* expression first begins to increase on day 2 of ALI epithelial differentiation (**Fig. 4c**). Therefore, we treated differentiating cultures at day 3 with IL13 (10 ng/μl) and profiled cells from days 5, 6 and 8 by scRNA-seq (**Fig. 4d**). Following profiling, we detected all the expected common cells as well as differentiating rare cells (**Extended Data Fig. 10a,b**).

In aggregate, we obtained a total of 572 rare cells from cultured control and IL-13 treated epithelial cells, and they partitioned into five major rare cell clusters (**Extended Data Fig. 10c,d**, **Extended Data Table 7**). We next individually clustered the control cells (**Fig. 4e-k**) and the cells that were exposed to IL13 (**Fig. 4l-r**). These included cycling POU2F3^+^ progenitor cells enriched for markers of active proliferation (**Fig. 4e-h, l-o**) and non-cycling POU2F3^+^ progenitor cells that express *ASCL2*, *POU2F3* and *FOXI1*, but not *ASCL3* (**Fig. 4e,k,l,r**). We also identified ASCL3^+^ pre-ionocytes (**Fig. 4e,k,l,r**); mature ionocytes expressing *BSND, CFTR, ATP6V1C2, ATP6V0A4,* and *CLCNKB* (**Fig. 4e,i,l,p**); and mature tuft cells expressing functional tuft cell genes (*IL25, CHAT, GNAT3, GNB3, GNG13, TAS1R1, TAS1R3, ALOX5, ALOX5AP*) (**Fig. 4e,j,l,q**).

Marker gene expression and RNA velocity analysis under normal conditions once again supports a model in which POU2F3^+^ progenitors give rise to ionocytes (**Fig. 4e,f**). Following IL13 treatment, POU2F3^+^ progenitors differentiate into mature tuft cells (**Fig. 4l,m**). Given the ability of POU2F3^+^ progenitor cells to give rise to both mature tuft and ionocyte lineages, we will hereafter refer to these cells as <u>T</u>uft Ionocyte <u>P</u>rogenitor cells (TIP cells). In ALI culture, many of these TIP cells proliferate, whereas pre-ionocytes, mature ionocytes, and mature tuft cells do not cycle (**Fig. 4e,g,h,l,n,o**). As is the case *in vivo*, the fraction of cycling TIP cells (49%) is once again higher than that of cycling canonical basal stem cells (23%) (**Extended Data Table 7)** . TIP cells express both ionocyte (*FOXI1*) and tuft cell (*ASCL2*, *POU2F3*) TFs, in both control and IL-13 conditions (**Fig. 4e,k,l,r**). During differentiation towards a mature tuft cell or ionocyte, the expression of TFs of the opposing lineage are suppressed (**Fig. 4e,k,l,r**). Of particular note, *SOX9* expression is induced in the tuft cell lineage and absent in the ionocytes, while conversely *ASCL3* expression presages ionocyte commitment and is absent in the mature tuft cells (**Fig. 4e,k,l,r**). Finally, this is followed by expression of functional tuft and ionocyte genes in the respective mature cells (**Fig. 4e,i,j,k,l,p,q,r**). In aggregate, TIP cells descendants proceed through an orderly process of differentiation associated with the sequential expression of distinct cell fate determining transcription factors based on the presence or absence of IL13 (**Fig. 4s**) Finally, prolonged IL-13 treatment of differentiating LAE ALI cultures decreased the number of ionocytes per ALI membrane and increased the number of mature tuft cells (**Fig. 4t**), presumably because cell turnover resulted in a loss of ionocytes that could not be replenished by TIP cells because TIP cells had been diverted towards a mature tuft cell fate. Similarly, small airway epithelial (SAE) ALI cultures treated with IL13 (10ng/μl) displayed increased mature tuft cell differentiation (**Extended Data Fig. 10e**). In aggregate, these findings point to a lineage tree in which basal stem cell-derived TIP cells can differentiate into either mature tuft cells or ionocytes depending on the signaling milieu (**Fig. 4u**).

### Type 17 cytokine signaling also induces tuft cell differentiation

Tuft cell differentiation is generally associated with Type 2 inflammation, but tuft cells are also found in human SARS-Cov2 pneumonia^19^ and in mouse models of influenza infection^49^, which are characterized by a combination of Type 1 and Type 17 inflammation. Indeed, the differentiation of murine influenza-induced tuft cells is independent of Type 2 cytokine signaling^50^. Furthermore, IL17A also induces a tuft-like gene signature in pancreatic cancer^51^. We therefore hypothesized that Type 17 cytokine signaling could also bias the lineage of TIP cells towards mature tuft cell differentiation since IL17A receptors (*IL17RA* and *IL17RC*) are expressed in TIP cells (**Extended Data Fig. 9e**).

Supporting our hypothesis, fully differentiated ALI cultures treated with IL17A (50 ng/μl) (**Fig. 5a**) displayed increased GNAT3^+^ tuft cell differentiation (number of cells per ALI membrane) (**Fig. 5b**). Moreover, continuous administration of IL17A for 21 days during ALI epithelial differentiation and maturation also increased tuft cell differentiation (**Fig 5c,d**), as was previously observed with the continuous administration of IL13 (**Fig. 4t**). Similarly, small airway epithelial (SAE) ALI cultures treated with IL17A (50ng/μl) also displayed increased mature tuft cell differentiation (**Extended Data Fig. 10e**). To our knowledge, this represents the first direct evidence for a role of Type 17 signaling in tuft cell formation (**Fig. 5e**).

### Mature tuft cells are present in human asthmatic airways

Since we have shown that mature tuft cells can be induced in airway epithelial cultures by type 2 cytokines, we hypothesized that airway epithelial cultures derived from asthmatic individuals should contain mature tuft cells, even though such cells have not heretofore been detected in the airways of asthmatic individuals *in situ*^3,46^.

To test our hypothesis, we isolated basal cells from the large airways of individuals that died during an asthma exacerbation and generated fully differentiated LAE ALI cultures (without added cytokines). These cultures contained GNAT3^+^DCLK1^+^ mature tuft cells (**Fig. 5f**). Moreover, there was a 2-4 fold increase in the number of GNAT3^+^ mature tuft cells (cells per airway membrane) in cultures grown from basal stem cells of asthmatic patients compared to those of individuals without a history of any lung disease (**Fig. 5g**). Interestingly, as the cultures are grown from basal cells, this suggests that the basal stem cells retain a memory of the distinct cytokine environment in the asthmatic airway^52–54^, consistent with the demonstration of allergic inflammatory memory in nasal polyp-derived basal stem cells^55^.

To verify whether this excess tuft cell differentiation *in vitro* reflects *in vivo* asthma biology, we performed whole mount staining of micro dissected airway using intact tissue from the one patient from whom intact tissue was available. Mature tuft cells were present in both the large and small airways of this patient (**Fig. 5h**), while none were detected in the large and small airways of normal human individuals. To our knowledge, this represents the first direct evidence for the existence of mature tuft cells *in situ* in asthmatic large and small airways.

## DISCUSSION

To systematically study rare epithelial cells of the human lung, we performed a deep sampling of cells from 8 normal donor lungs. Querying this very large dataset, alongside the first version of the Human Lung Cell Atlas^16^, allowed us to amass a sufficient number of rare cells for detailed computational analysis. We then amassed a complementary dataset of rare cell scRNAseq profiles obtained from *in vitro* epithelial culture experiments in which the rare cells had first been enriched through cell sorting. By combining this dataset with our *in vivo* dataset, we described the identity and physiologic regulation of the TIP cell, a new replicative rare cell progenitor that has the potential to differentiate into either mature tuft cells or ionocytes. Indeed, absent the ability to perform prospective clonal assays (as was used to identify rare progenitor cells in hematopoiesis) there is currently no other methodology for delineating the mechanisms governing the behavior of exceedingly rare cells from solid tissues.

TIP cells were named following the simple convention for the naming of hematopoietic progenitors, based on their spectrum of differentiation. Mechanistically, TIP cell bipotency is associated with the dual expression of TFs of both the tuft and ionocyte lineage. Prior studies that had cataloged these rare progenitors had variously annotated them as “tuft-like”^1^, “tuft”^16^, or “undefined rare”^5^ cells, but, in fact, the default pathway of differentiation of TIP cells produces ionocytes. Indeed, *bona fide* mature tuft cells are almost entirely absent in the normal human airway. Remarkably, 18.35% of TIP cells are replicating *in vivo*, exceeding the cycling fraction of basal stem cells. Interestingly, this high rate of turnover suggests the possibility that ionocytes, the default products of TIP cell differentiation, have a short half-life and therefore require constant replenishment.

Previously identified more proximal tracheal progenitor cells, termed “tuft-like” cells, also expressed genes associated with both tuft cell and ionocyte differentiation, but it was suggested that these “tuft-like” cells could serve as progenitors for ionocytes and neuroendocrine (NE) cells, despite the fact that these tracheal tuft-like cells did not express markers of NE cell differentiation^1^. However, the lack of sufficient numbers of cells did not permit the identification of transitional cells that could directly establish lineage relationships. Similarly, the lineage relationship of these tuft-like cells to mature tuft cells could not be assessed given the absence of mature tuft cells. Of further interest, the developing ferret possesses a tripotent rare cell progenitor population, but this progenitor cell population could not be identified in the adult ferret airway ^12^. Based on these and our observations in aggregate, we speculate that it is possible that TIP cells may demonstrate tripotency and give rise to NE cells under the appropriate conditions. However, at homeostasis, we find no evidence of such transitional cells despite the depth of our cell atlas and the identification of many mature NE cells. Perhaps in the setting of rare human genetic disorders characterized by an excess of NE cells, such as NEHI (Neuroendocrine hyperplasia of infancy) ^56^, TIP cells can be pathologically diverted to produce excess NE cells. Alternatively, bronchial solitary NE cells may arise from a separate distinct rare cell progenitor population, as seen in the developing human lower airway ^57^ or could be sourced directly from basal cells as we have previously shown^56^.

Our work suggests that the behavior of TIP cells is altered in human airway disease. First, we showed that TIP cell lineage preference is modulated by Type 2 and Type 17 cytokines, which are both characteristically expressed in inflammatory airway diseases. Furthermore, we report the first *in situ* identification of human mature tuft cells in asthma. We speculate that excess sentinel tuft cells, and attendant increased IL25 and leukotriene secretion, may create a milieu in which an innocuous stimulus triggers pathologic type 2 inflammation, which could, in turn, create a cycle in which tuft cell differentiation is further enhanced. Since excessive type 2 signaling is characteristic of allergic inflammation and excessive type 17 signaling is associated with neutrophilic asthma, these findings nominate the TIP cell as a potential target for therapeutic interventions in settings where tuft cell expansion might be promoting pathogenic inflammatory responses. Additionally, the finding that asthmatic basal cells, the parents of TIP cells, produce mature tuft cells when cultured *ex vivo* suggests that they retain a memory of the distinct cytokine environment of the asthmatic airway ^52–54^ that results in a biasing of TIP cell differentiation towards the mature tuft cell fate, consistent with the demonstration of allergic inflammatory memory in nasal polyp-derived basal stem cells ^55^.

With regard to the distribution of ionocytes, direct whole mount tissue analysis of dissected large and small airways identified the presence of CFTR-rich distal ionocytes in indistinguishable numbers along the proximodistal axis of the human airway tree. Recently, experiments in which ionocytes were genetically ablated in the ferret led to aberrant airway surface physiology mirroring the abnormalities seen in CF ^12^ while the loss of human ionocytes led to abnormalities of ion transport *in vitro* ^14^. Taken together, this suggests that human ionocytes likely play a role in the pathogenesis of both large and small airways disease in CF patients. Indeed, the prior assertion that CFTR-harboring secretory cell dysfunction is the primary cause of small airways disease in CF has been called into question since it has recently been suggested that the prior inability to identify ionocytes in the distal airway may be due to their fragility and the resulting spurious expression of CFTR in secretory cells ^12,24^. Given our demonstration of the presence of distal human airway ionocytes, it may be crucial to design gene therapy strategies that target both the large and small airways to ensure the production of corrected ionocytes in both anatomic compartments of the airway tree. Along these lines, our scATAC-seq studies identified transcriptional regulators that control rare cell lineages and likely determining the course of differentiation of TIP cells into pre-ionocytes and then ionocytes, offering potential candidate TFs for reprogramming cells into ionocytes. Finally, the use of *ex vivo* culture systems coupled to our rare epithelial cell enrichment strategies should provide a methodology for the further mechanistic interrogation of rare cell pathobiology in a host of distinct airway diseases. Taken together, our findings suggest that diseases associated with airway inflammation may more generally display a skewing of normal tuft and ionocyte distributions, potentially resulting in a disruption of normal airway physiology, mucociliary transport, and host defense. More generally, our study highlights the power of very deep sampling of primary human tissues. This is especially important since rare cells are now being increasingly recognized to have outsized roles in physiology and pathophysiology. Indeed, we speculate that the diversity and potency of rare cell progenitors and their respective lineages will vary based on species ^1,7,12^, by developmental stage ^12,13^, and by anatomic location ^16^. Finally, as a corollary, our studies suggest that the lineage of rare cell progenitors will be biased in disease states which may in turn serve as a substrate that promotes airway dysfunction and disease.

## METHODS

### Human lung tissue

Human lung tissue was received from New England Donor Services under the Massachusetts General Hospital approved Institutional Review Board protocol [Protocol # 2010P001354]. Tissue was collected from individuals without any reported history of lung disease, smoking or drug use.

### Human lung tissue dissection, dissociation and cell enrichment sorting

Large airways were manually dissected and washed with HBSS media (Sigma) containing Y-27632 (5 μM) inhibitor in order to remove any mucus and debris. Single cells were dissociated in HBSS media containing pronase (1 mg/ml), DNAse (1 unit/ml) and Y-27632 inhibitor (5 μM) on a 37°C shaker for 30-60 minutes. The dissociated epithelial layer was collected by centrifugation. Cell pellets were treated with TrypLE™ Express solution (ThermoFisher Scientific) for 5 minutes at 37°C with rotation. Cell suspensions were collected by centrifugation and treated with ACK lysis buffer (ThermoFisher Scientific A1049201) for 1 minute on ice. Cell pellets were resuspended in phosphate buffered saline (PBS) with 10% FBS and Y-27632 inhibitor (5 μM) and then sequentially filtered through 100µM and 40µM filters. Cell viability was assessed using a hemocytometer and single cell suspensions were placed on ice for further experiments. To further enrich large airway rare epithelial cells we used FACS sorting. We stained cells with anti-human CD45–BV605 (1:100; BioLegend) or anti-human CD45–BV421 (1:100; BioLegend), anti-human CD31–BV421 (1:100; BioLegend), anti-human EPCAM–APC or EPCAM–PE (1:100; BioLegend), anti-human CD117–FITC or CD117–APC-Cy7 (1:100; BioLegend) for 20 minutes at room temperature. Cells were washed and suspended in PBS containing 10% FBS and Y-27632 inhibitor (5 μM) with added DAPI solution to assess cell viability. After washing, cells were sorted on a FACS sorter and analysis was performed using FlowJo software.

Distal lung lobe regions were manually dissected, avoiding all airways larger than 2mm, and washed with HBSS containing Y-27632 inhibitor (5 μM) to remove any mucus and debris. Single cells were dissociated in HBSS media (Sigma, 55021C) containing collagenase (225 units/ml), dispase (2.5 units/ml), elastase (2 units/ml), pronase (1 mg/ml), DNAse (1 unit/ml) and Y-27632 inhibitor (5 μM). Tissue fragments were injected with the above dissociation solution multiple times with a syringe and then divided into smaller pieces that were placed in a 37°C shaker for 60 minutes. Cell suspensions were treated with ACK lysis buffer (ThermoFisher Scientific A1049201) for 2 min on ice to remove red blood cells. Cell pellets were resuspended in PBS with 10% FBS and Y-27632 inhibitor (5 μM) and sequentially filtered through 100µM and 40µM filters. Cell viability was assessed using a hemocytometer and single cell suspensions were placed on ice for further experiments. Additionally, in order to enrich for rare cell types in distal lung lobe regions, we used sequential magnetic cell separation using MACS cell separation columns on three human lung samples (Hu37, Hu49 and Hu62). Specifically, for Hu37 and Hu49 distal lung lobe samples, we used human CD45 MicroBeads (Miltenyi Biotec catalog number 130-045-801) to enrich immune cells in the CD45 positive fraction and further loaded the CD45 negative fraction onto CD31 MicroBeads (Miltenyi Biotec catalog number 130-091-935). We collected the CD31 positive fraction to enrich endothelial cells and also collected and loaded the CD31 negative fraction. For Hu62, distal lung lobe samples, we used human CD45 MicroBeads (Miltenyi Biotec catalog number 130-045-801) to enrich immune cells in the CD45 positive fraction and further loaded the CD45 negative fraction onto CD31 MicroBeads (Miltenyi Biotech catalog number 130-091-935). We collected the CD31 positive fraction to enrich endothelial cells and the CD31 negative fraction was then loaded onto human anti-IgM MicroBeads conjugated to anti HTII-280 antibody (Terrace Biotech catalog number TB-27AHT2-280) to enrich AT2 cells. We collected the HTII-280 negative fraction to increase the number of captured small airway epithelial cells.

### Dissection of large and small airways and whole mount staining

Large and small airway samples were handled identically. Within 24 hours after explanting the lungs, airways were dissected at various indicated areas in the right and left mainstem bronchi and the right and left distal airways (<2mm in diameter; as in **Extended Data Fig. 7**). Tissue explants were then fixed in 4% PFA for 1 hour and washed in PBS for 1 hour. Tissues were preserved in PBS until staining. Explants were then permeabilized in PBS-0.3% TritonX-100 (PBST) for 30 minutes. Samples were stained with primary antibody at 4°C overnight, diluted in 1% BSA-0.3% PBST. The following antibodies and dilutions were used:

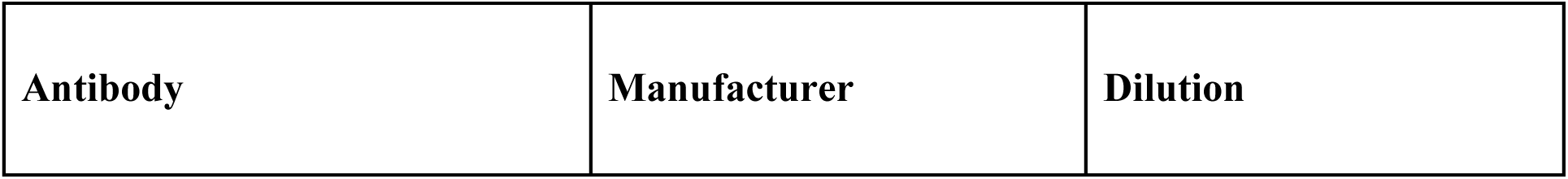
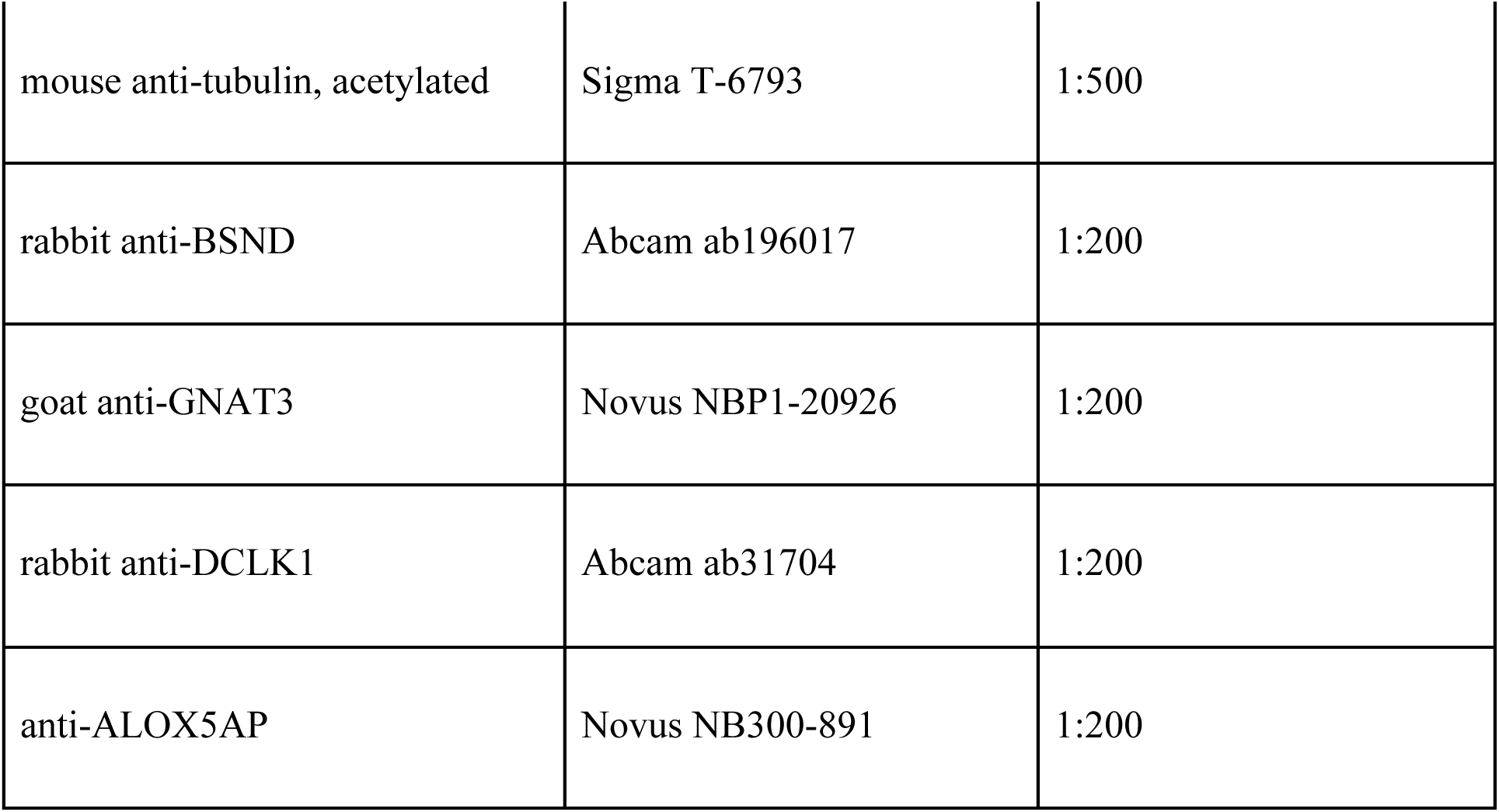

| **Antibody** | **Manufacturer** | **Dilution** |
| --- | --- | --- |
| mouse anti-tubulin, acetylated | Sigma T-6793 | 1:500 |
| rabbit anti-BSND | Abcam ab196017 | 1:200 |
| goat anti-GNAT3 | Novus NBP1-20926 | 1:200 |
| rabbit anti-DCLK1 | Abcam ab31704 | 1:200 |
| anti-ALOX5AP | Novus NB300-891 | 1:200 |

For each donor, 3 large airway and 3 small airway samples were obtained. From these, a number of airways (each 0.06480mm^2^ surface area region) were obtained for each donor based on the size of tissue dissected, as follows: HU66: 8 large airway regions and 9 small airway regions; HU67: 10 (large) and 17 (small); HU68: 8 (large) and 7 (small). Each explant was mounted onto an O-ring (McMaster Carr 1283N264) fixed into a 60mm dish (Cellstar 628160) with Sylgard 184 silicone elastomer (EMS Catalog #24236-10). Each region was imaged with a 25X objective and cells were counted on a maximal intensity projection. Only regions in which there was concomitant staining for acetylated tubulin were included to ensure imaging of the airway epithelial layer. For BSND staining, 1.68mm^2^ of large airway and 2.14mm^2^ of small airway were imaged in total. ANOVA (Sidak’s multiple comparisons) was done. Each dot represents individual fields imaged on respective donors.

### Air liquid interface cultures

Cell were dissociated from large airways or distal lung lobe regions and expanded in basal cell media containing PneumaCult™-Ex Plus medium (Stem Cell Tech) containing 0.5 μM DMH-1 (Tocris #4126), 1μM CHIR 99021 (Tocris #4423), 1μM A-8301 (Tocris #2939), 5μM Y-27632 (Selleckbio #S1049) and Primocin 500x (Invivogen #ant-pm-2). To initiate air–liquid interface (ALI) cultures, basal cells were seeded onto transwell membranes with basal cell media in the upper and lower chambers. After reaching confluence, media was removed and replaced with PneumaCult™-ALI Medium (Stem Cell Tech #05001) in both the upper and lower chambers. After 24 hours, media was removed from the upper chamber to begin basal cell differentiation for the indicated number of days. ALI cultures were treated with either 10 or 20 ng/ml of recombinant human IL13 (Peprotech), 10 ng/ml of recombinant human IL4 (Peprotech), 10 ng/ml of recombinant human IL5 (Peprotech), or 50 ng/ml of recombinant human IL17A (Peprotech) as indicated (all diluted in PneumaCult-ALI Medium). We initially used 20 ng/ml of IL13 in ALI cultures as previously reported^7,58^, but subsequently found that a lower concentration of 10 ng/ml IL13 was sufficient to induce mature tuft cell and goblet cell differentiation, so this concentration was employed in the data presented in Figure 4. Control cultures received an equal volume of PBS or DMSO.

### Staining and quantification of cells in Air liquid interface cultures

The luminal side of the ALI membranes were washed with 200ul of PBS or 10mM DTT in PBS for 1 minute followed by an additional wash with PBS. ALI membranes were fixed in 4% PFA for 15 minutes and washed in PBS for 5 minutes. ALI membranes were preserved in PBS until staining. ALI membranes were permeabilized in PBS-0.3% TritonX-100 (PBST) for 60 minutes.

Samples were stained with primary antibody at 4°C overnight, diluted in 3% BSA 0.5% Tween20 5% Donkey Serum in PBS. Samples were stained with secondary antibodies at room temperature for 1 hour, diluted in 3%BSA 0.5%Tween20 5% Donkey Serum in PBS. The primary and secondary antibodies used and their concentrations are listed in the table below.

| **Antibody** | **Manufacturer** | **Dilution** |
| --- | --- | --- |
| rabbit anti-BSND | Abcam ab196017 | 1:200 |
| goat anti-GNAT3 | Novus NBP1-20926 | 1:100 |
| rabbit anti-DCLK1 | Abcam ab31704 | 1:100 |
| goat anti-ALOX5AP | Novus NB300-891 | 1:100 |
| Alexa Fluor 488 donkey anti-rabbit IgG | Thermo Fisher Scientific, A-21206 | 1:500 |
| Alexa Fluor 594 donkey anti-rabbit IgG | Thermo Fisher Scientific, A-21207 | 1:500 |
| Alexa Fluor 647 donkey anti-rabbit IgG | Thermo Fisher Scientific, A-31573 | 1:500 |
| Alexa Fluor 488 donkey anti-goat IgG | Thermo Fisher Scientific, A-11055 | 1:500 |
| Alexa Fluor 594 donkey anti-goat IgG | Thermo Fisher Scientific, A-21447 | 1:500 |
| Alexa Fluor 647 donkey anti-goat IgG | Thermo Fisher Scientific, A-11058 | 1:500 |

The ALI membranes were then mounted using a mounting medium containing DAPI. Confocal images were obtained using an Olympus FV10i confocal laser-scanning microscope with a 10× objective and the images were processed using Image J and cells from the ALI membrane were manually counted. Statistical analysis for Figure 1i was a 2 tailed unpaired T test. Each dot represent counts on ALI membrane for individual donors. Statistical analysis for Extended data Figure 6a one way ANOVA. P=0.0001.

### Generation of GCaMP6s cell line and live visualization of calcium signals

Human airway basal cell lines were transfected with lentivirus carrying CMV-GCaMP6s and a puromycin selection marker. Positively transfected cells were selected with 1μg/ml puromycin for 3 consecutive days. Basal cells were then grown in air liquid interface cultures as described above. After maturation, ALI membranes were cut from transwell and mounted on a glass bottom imaging chamber (Warner Instruments #64-0228). GCaMP signals were visualized using an Olympus IX81 inverted epifluorescence microscope fitted with a blue excitation filter.

### ALI membrane cell dissociation and FACS

ALI membranes were cut and washed twice with PBS. Membranes were incubated in TrypLE™ Express solution (ThermoFisher Scientific) and incubated at 37 °C for 5 minutes with rotation. Dissociation was stopped with PBS containing 10% FBS and Y-27632 inhibitor (5 μM).

Dissociated cells were passed through a 40µM filter. Cells were then pelleted and stained with anti-human CD45–BV605 (1:100; BioLegend), anti-human CD31–BV421 (1:100; BioLegend), anti-human EPCAM–APC or EPCAM–PE (1:100; BioLegend), anti-human CD117–FITC or CD117–APC-Cy7 (1:100; BioLegend), and anti-human CD56–BV711 (1:100; BioLegend) for 20 minutes at room temperature. Cells were washed and suspended in PBS containing 10% FBS and Y-27632 inhibitor (5 μM) with added DAPI solution to assess cell viability. After washing, cells were sorted on a BD FACS Aria (BD Biosciences) using FACS Diva software and analysis was performed using FlowJo software.

### scRNA-seq library generation and sequencing

Droplet-based scRNA-Seq 10x Genomics V2 (donor samples from Hu28, Hu30, Hu32, Hu37, Hu39, and Hu49) or V3 (donor samples from Hu52 and Hu62) 3’ gene expression technology was used according to manufacturer’s recommendations. In short, single cells were washed with 0.4% BSA in PBS, and loaded onto a Chromium single-cell 3′ Chip. Single cells were partitioned into droplets with gel beads in the Chromium Controller. After emulsions were formed, barcoded reverse transcription of RNA took place. This was followed by cDNA amplification, fragmentation and adapter and sample index attachment. Libraries were pooled together and sequenced on an Illumina NextSeq 500, or an Illumina Nova-Seq, with paired end reads.

### Pre-processing of human lung scRNA-seq data

Raw 3’ scRNAseq sequencing BCL files were demultiplexed using the CellRanger pipeline (10X Genomics) version 2.1.0. *‘mkfastq’* command using default parameters to generate *fastq* files followed by *‘count’* command to align the reads to the human GRCh38 reference genome.

### Quality control and data analysis of human lung scRNA-seq datasets

The resulting raw barcode, gene, and count matrices from individual samples were individually loaded in R(4.0.5) for downstream analysis using Seurat^59^ version 4.0.6 with the *read 10X* function. We removed low quality cells with fewer than 1000 unique molecular identifiers (UMIs), fewer than 400 detected genes or greater than 25% mitochondrial genes from all downstream analysis. The remaining cells were ‘LogNormalized’ using NormalizeData() function and 2000 variable genes were identified using the default ‘vst’ method in the ‘FindVariableFeatures’ function. The normalized data was scaled using the ScaleData function, which was then used to perform principal component (PCA) analysis on the variable gene expression space. Next, dimensionality reduction of the first 20 principal components was performed using Uniform Manifold Approximation and Projection (UMAP) for visualization and annotation of each donor lung scRNA-seq data prior to performing donor-wise integration.

### Cell clustering and annotation of human lung scRNA-seq

Cell type identification was performed by assigning cell clusters to cell subsets at different resolutions. The ‘FindNeighbors’ function was applied to each donor’s normalized gene-expression data matrix to calculate the pairwise distances between the cells and construct KNN and shared nearest neighbor (SNN) graphs. Then de novo clustering was performed using the Louvain algorithm at different resolutions (0.2, 0.4, 0.8, 1.2, 2.4 and 5) on the SNN graph space. For each cluster, we used the findAllMarkers() function to identify the positive markers for that cluster with a minimum fraction of 0.25, log2 fold-change of 0.25, where we used the Wilcoxon rank-sum test for finding upregulated differentially expressed genes (DEGs) for each cluster and computed Bonferroni corrected p-values. We overlapped de novo discovered DEGs with well-established cell type markers to perform high-level cell type annotation in an iterative and semi-supervised way to assign clusters to broad classes based on consistent expression of known markers in a given cluster. We used the following markers for broad cell class annotation: epithelial (*EPCAM, KRT8, KRT18, TP63, S100A2*), mesenchymal (*COL1A1, COL1A2, DCN, FBLN1*), myeloid (*LYZ, C1QA, C1QB, APOC1*), T (*CD3D, CD3E, GZMK*) & NK (*GNLY, GZMB, KLRD1*), B/plasma (*CD79A, MS4A1, MZB1, JCHAIN, IGHG1*), mast (*TPSAB1, TPSB2, CPA3*), and endothelial (*VWF, PECAM1, CLDN5*). We then subset cells within each of the broad classes and further subset samples into two groups (proximal and distal) based on whether they were acquired from the large airways (proximal) or parenchyma (distal). To annotate cell subsets within each broad cell class from either proximal or distal region samples, the whole process of log normalization, selection of “variable” genes, scaling, dimensionality reduction (PCA/UMAP), clustering at different resolutions, and finding DEGs was repeated, as described above. Doublet enriched clusters were identified as clusters expressing canonical markers for two cell types and containing higher numbers of genes per UMI (genes/UMI) than expected from each individual cell type. These doublets were removed from downstream analysis.

### Data integration of human lung scRNA-seq

There was a noticeable batch effect in our human lung data set primarily driven by using either V2 or V3 chemistry 3’ gene expression kits for different donors, which required robust data integration. We chose to integrate the data across donors using the anchor-based approach as implemented in Seurat v3^59^. Prior to integration, count matrices for individual donors were normalized and mitochondrial gene content per cell was regressed out using the SCTransform() function. Next, we selected 3000 integration features via the SelectIntegrationFeatures() function followed by donor-wise integration using PrepSCTIntegration(), FindIntegrationAnchors(), and IntegrateData() function respectively. Post-integration, the resulting integrated Seurat objects were subjected to downstream analysis, as described above, including data log normalization, selection of variable features, data scaling, dimensionality reduction (PCA/UMAP), clustering, finding DEGs, and cell annotation.

### scRNA-seq cell cycle analysis

To identify cycling cells, we used the Seurat CellCycleScoring() function to compute, for each cell, the enrichment score for the expression of genes linked to either G1/S or G2/M phase of the cell cycle^60^. Cells with either a G1/S- or G2/M-score greater than 0.1 were classified as cycling and all other cells were considered non-cycling (**Fig. 2d**). Finally, the resulting cycling status or cell cycle phase of cells were visualized using Seurat’s DimPlot() function for different UMAP embedding. To quantify cycling cell stages in the ALI scRNA-seq data, raw FASTQ files were aligned to the human reference build GRCh38 using CellRanger 7.0.1 ‘count’ function, input into Scanpy, and scored with the integrated list of cell cycle genes used in the approach implemented in Seurat^60^ via the score_genes_cell_cycle() function with the default n_bins=25.

### scRNA-seq module score analysis and detection of mature tuft cells

Sub-clustering of epithelial cells in our scRNA-seq data allowed us to identify clusters containing ionocytes, neuroendocrine (NE) cells, and TIP cells, but not mature tuft cells. To detect mature tuft cells in our data, we utilized published scRNA-seq data of rare lung epithelial cells ^5^ and identified the top ten differentially expressed genes for ionocytes, NE, TIP, and mature tuft cells (**Extended Data Table 2**). Signature scores for rare epithelial cells were assigned per cell using the AddModuleScore() function in Seurat v4, which compares the average expression of genes in an input gene set to the aggregate expression of control sets of genes randomly selected from the same expression level bins as the input gene set^60^.

### HLCA rare epithelial cell analysis

To characterize the number and proportion of rare lung epithelial cells in other published data sets, we subset the rare epithelial cells in the main HLCA v1 data set using Seurat v4. We log normalized the data using the NormalizeData() function and identified 2000 variable features using the FindVariableFeatures() function. Next, we performed data scaling, PCA, batch correction using Harmony (v1.0), and UMAP dimensionality reduction to visualize the data in two dimensions (**Extended Data Fig. 4a-c**). Finally, we scored all cells for rare lung epithelial cell signatures (**Extended Data Table 2**) using the AddModuleScore() function and assigned clusters that were specifically enriched for one of the 4 rare cell type signatures of the corresponding rare cell identity.

### RNA velocity analysis of scRNA-Seq

Aligned scRNA-seq reads were further processed using the Velocyto 0.17^61^ read counting pipeline in Python 3.6.0. The GRCh38 expressed repeat mask was used. The output loom file was processed using Scanpy 1.6.0 with leidenalg 0.8.3. For *in vivo* human lung scRNA-seq data, quality control and integrated UMAP plots were obtained as described above. For ALI scRNA-seq data, high quality cells were retained, those that had >200 genes detected, <15% mitochondrial read count, and >.45 doublet prediction score from scrublet (2020 version) (automatic detection threshold, manually verified). scVelo 0.2.2 was used to predict RNA velocity with the following parameters: n_neighbors=20 (for full data set) and n_neighbors=5 (for subset computation). Stochastic, deterministic, and dynamical modeling were all tested, and quality control plots of known key marker genes were plotted to select stochastic estimation as the final modeling method.

### Single cell ATAC-Seq (scATACseq) library generation and sequencing

scATAC-seq was performed from primary carina and subpleural parenchymal tissue of one donor (Hu62). Libraries were generated using the 10x Chromium Controller and the Chromium Single Cell ATAC Library & Gel Bead Kit (1000111) according to the manufacturer’s instructions (CG000169-Rev C; CG000168-Rev B) with the following modifications in cell handling and processing. Briefly, human lung primary cells were processed in 1.5 ml DNA LoBind tubes (Eppendorf), washed in PBS via centrifugation at 400*g* for 5 min at 4 °C and lysed for 3 min on ice before washing via centrifugation at 500*g* for 5 min at 4 °C. Supernatant was discarded and lysed cells were diluted in 1× diluted nucleus buffer (10x Genomics) before counting using trypan blue and a Countess II FL Automated Cell Counter to validate lysis. If large cell clumps were observed, a 40-µm Flowmi cell strainer was used before the tagmentation reaction, followed by Gel Bead-In-Emulsion generation and linear PCR as described in the manufacturer’s protocol (10x Genomics). After breaking the emulsion, barcoded tagmented DNA was purified and further amplified to enable sample indexing and enrichment of scATAC-seq libraries. Final libraries were quantified using a Qubit dsDNA HS Assay kit (Invitrogen) and a High Sensitivity DNA chip run on a Bioanalyzer 2100 system (Agilent). All libraries were sequenced using NextSeq High Output Cartridge kits and a NextSeq 500 sequencer (Illumina), and 10x scATAC-seq libraries were characterized by paired-end sequencing (2 × 72 cycles).

### scATACseq data analysis

Fastq files were aligned to human genome reference hg19 using cellranger-atac (v1.2). Fragment files were parsed with ArchR (v1.0.2) and initial QC on cells was applied based on sequencing quality and depth, discarding cells with < 1,000 fragments or a transcription start site (TSS) enrichment score < 4. Dimensionality reduction and clustering was applied on cells passing filters using the standard ArchR analysis workflow. Cell type annotation of scATAC-seq data was guided using cell annotation labels from scRNA-seq data which were transferred to the scATAC-seq data using the function addGeneIntegrationMatrix. De novo marker discovery was run with the function getMarkerFeatures using the GeneScoreMatrix assay. Final scATAC-seq cell annotation was performed by assigning the de novo discovered clusters to cell types predicted by the RNA or to cell types associated with the most enriched scATAC-seq features in that cluster when no analog cell type was found in the RNA data. TF activity was inferred using ChromVAR (v1.16) implemented by ArchR via addMotifAnnotations, addBgdPeaks and addDeviationsMatrix functions. For visualization, ArchR built-in plotting functions and ComplexHeatmap were used (v2.10).

### Data and code availability

The single-cell RNAseq and scATACseq data from donor lungs was deposited in the Sequence Read Archive (SRA) with accession number SUB13969695. The ALI scRNAseq data was deposited at GEO with accession numbers GSE240168 (reviewer token:wlmbececftczjkp). Code and additional information required to reanalyze the data reported in this paper is available from the corresponding authors upon request.

## Extended Data Figure Legends

**Extended Data Figure 1.**
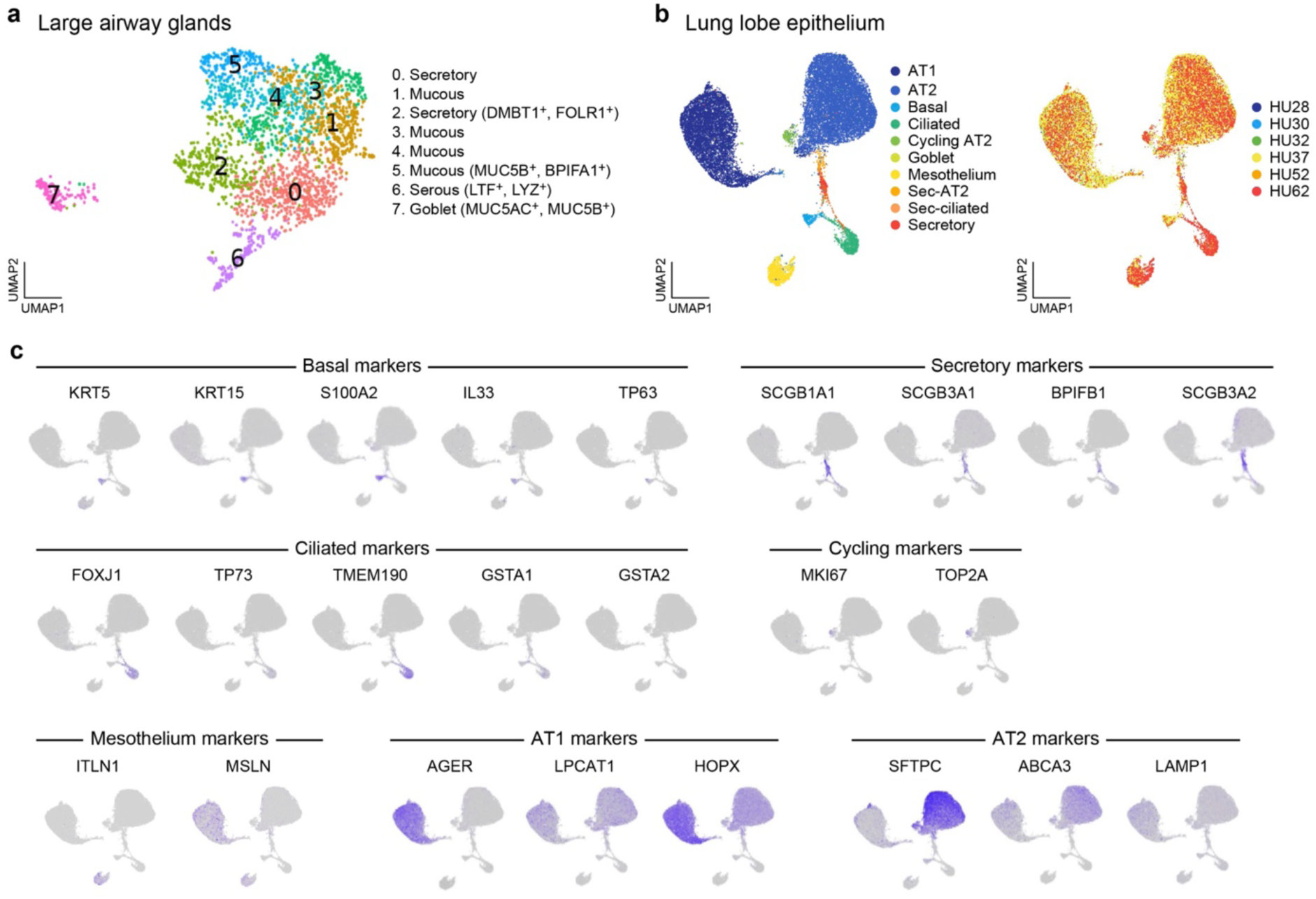
Large airway gland and alveolar epithelial cell subsets from large airways and lung lobes. **a.** Large airway gland cell subsets. UMAP embedding of gland cell scRNA-seq profiles (dots) from the large airways, colored by clustering and annotated post hoc. **b-c.** Lung lobe epithelial cell subsets, UMAP embedding of epithelial cell scRNA-seq profiles (dots) from the lung lobe, colored by cell annotation (b, left), donor lung (b, right), or gene markers of each subset (c).

**Extended Data Figure 2.**
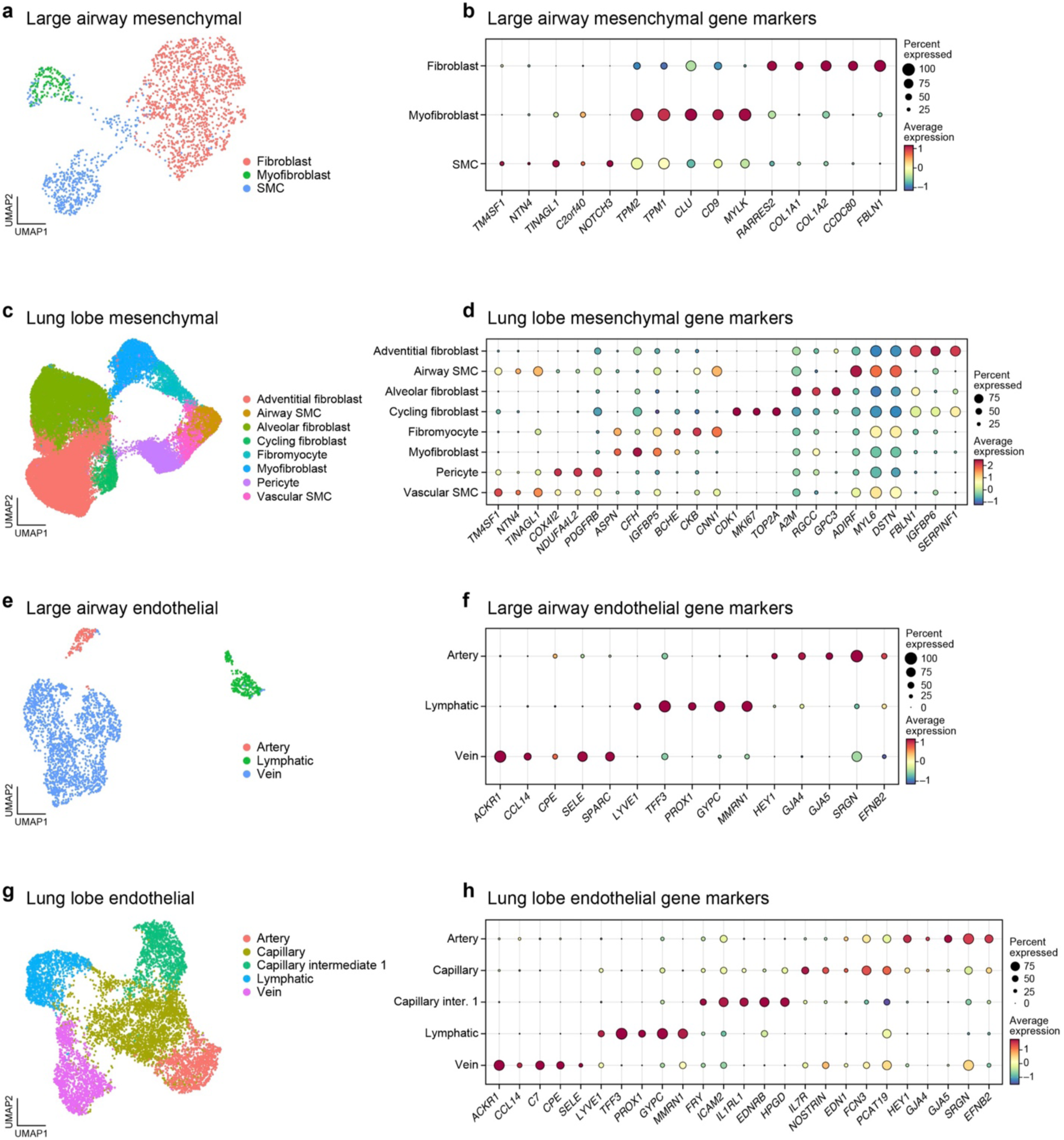
Mesenchymal and endothelial cell subsets from large airways and lung lobes. **a-d.** Mesenchymal cell subsets. UMAP embedding of mesenchymal scRNA-seq profiles (dots) from large airways (a) or lung lobe (c) regions colored by HLCA ^16^ annotations. **b,d.** Mean expression (dot color) and percent of expressing cells (dot size) gene markers (columns) of mesenchymal cell clusters from large airway (b) or lung lobe (d) regions. **e-h.** Endothelial cell subsets. UMAP embedding of endothelial scRNA-seq profiles (dots) from large airways (e) or lung lobe (g) regions colored by HLCA ^16^ annotations. **f,h.** Mean expression (dot color) and percent of expressing cells (dot size) gene markers (columns) of endothelial cell clusters (rows) from large airway (f) or lung lobe (h) regions. SMC - smooth muscle cells and Inter. - Intermediate.

**Extended Data Figure 3.**
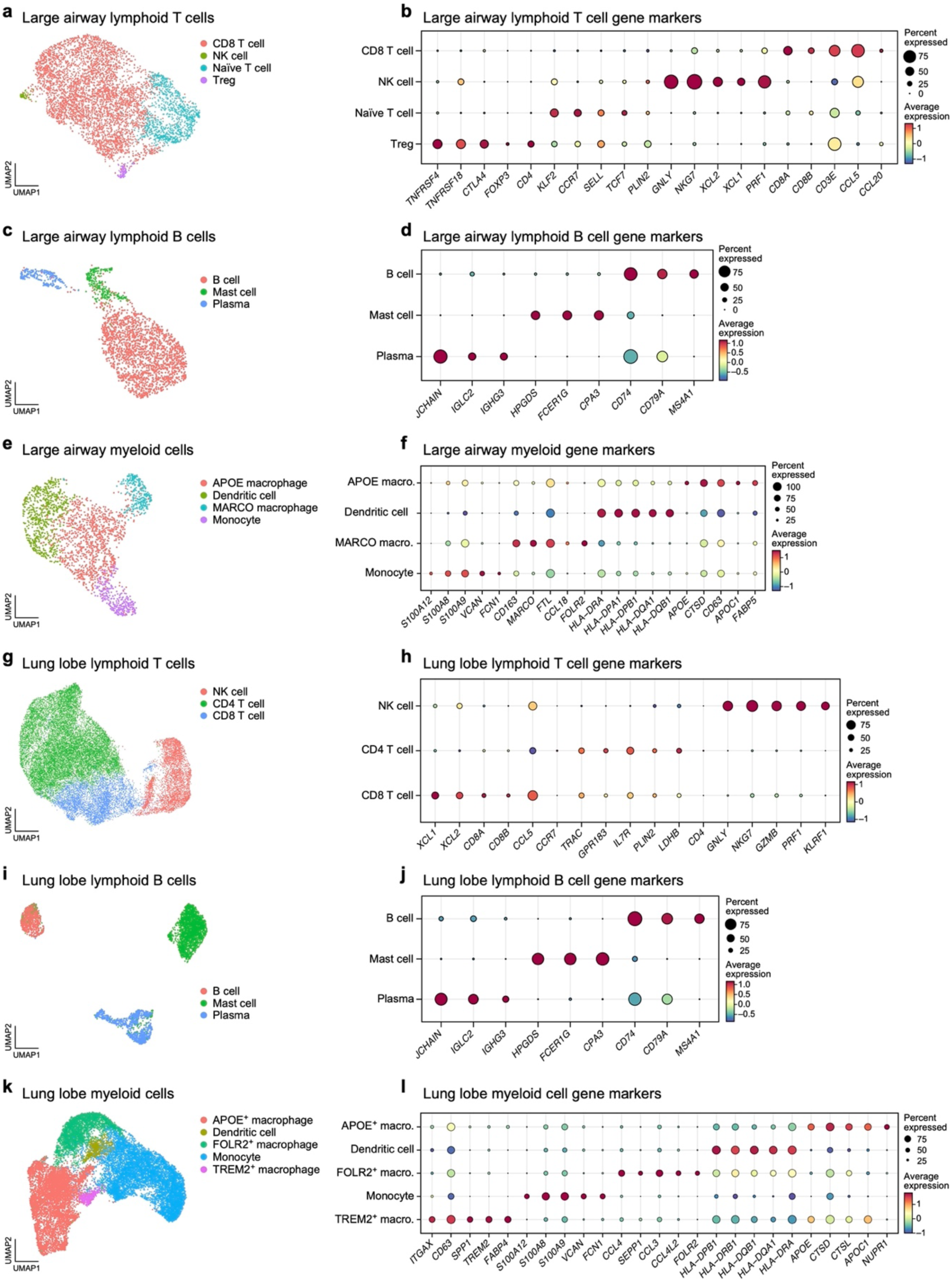
Immune cell subsets from large airways and lung lobes. **a,c,e,g,i,k.** UMAP embedding of scRNA-seq profiles (dots) of lymphoid T/NK (a), lymphoid B (c), and myeloid (e) cells from the large airways, and lymphoid T/NK (g), lymphoid B (i), and myeloid (k) cells from the lung lobes, colored by cell annotation. **b,d,f,h,j,l.** Mean expression (dot color) and percent of expressing cells (dot size) gene markers (columns) of lymphoid T/NK (b, rows), lymphoid B (d, rows), and myeloid (f, rows) cells from the large airways, and lymphoid T/NK (h, rows), lymphoid B (j, rows), and myeloid (l, rows) cells from the lung lobes. Macro.- Macrophages

**Extended Data Figure 4.**
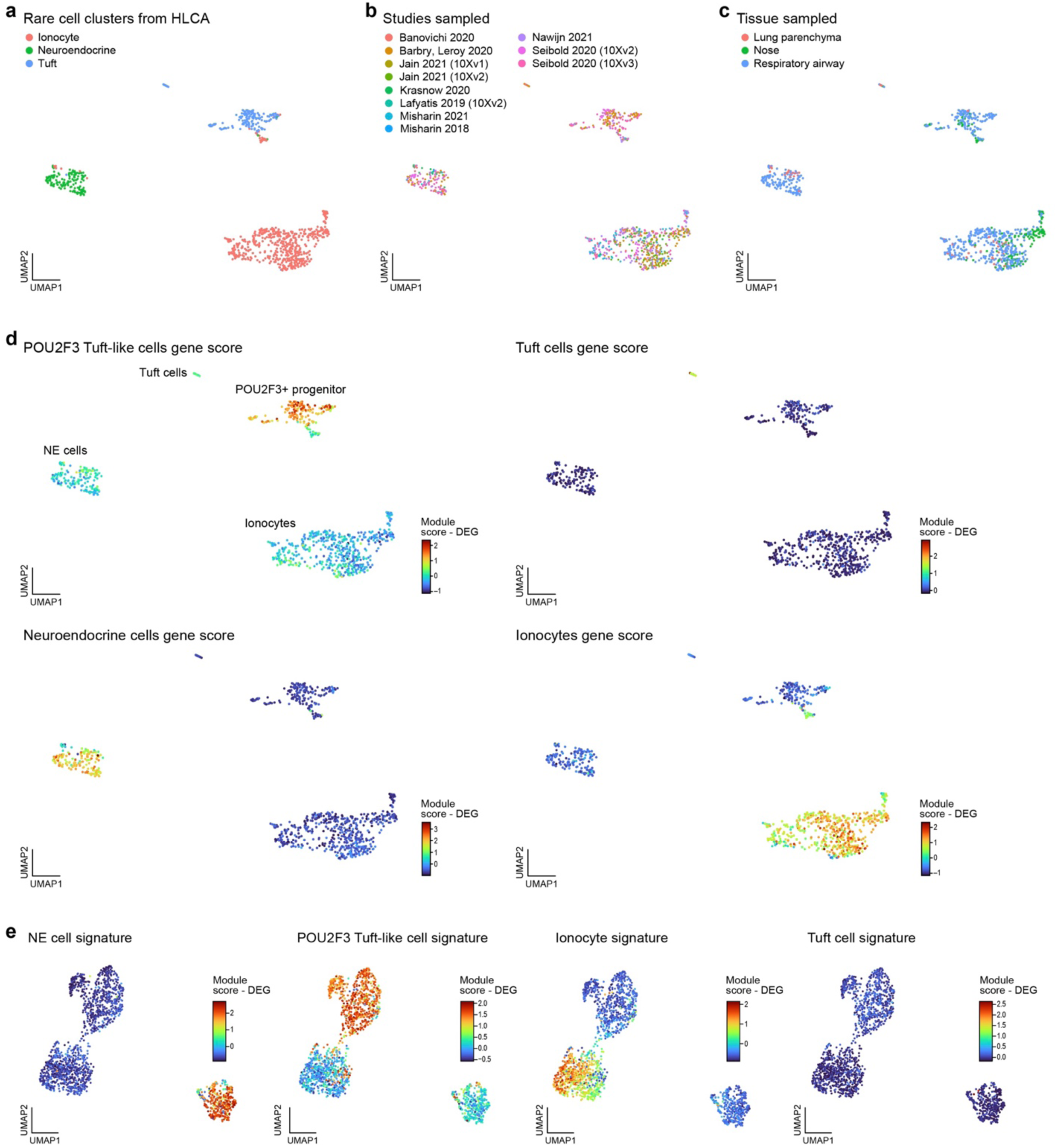
POU2F3^+^ “tuft-like” cells and mature tuft cells in the HLCA dataset and our deep lung atlas. **a-d.** POU2F3^+^ “tuft-like” cells are distinct from mature tuft cells in the HLCA dataset. UMAP embedding of rare cell profiles (dots) from HLCA v1.0 ^16^, colored by their HLCA annotations (a), the source studies (b), originating lung regions (c) and the by their re-annotation to POU2F3^+^ tuft-like cells, mature tuft cells, ionocytes and neuroendocrine cells based on gene module scores. **e.** POU2F3^+^ “tuft-like” cells are distinct from mature tuft cells. UMAP embedding of rare cell profiles (dots) from our deep lung atlas colored by annotation to neuroendocrine, POU2F3^+^ “tuft-like” cells, ionocytes and mature tuft cells based on gene module scores. DEG - differentially expressed genes

**Extended Data Figure 5.**
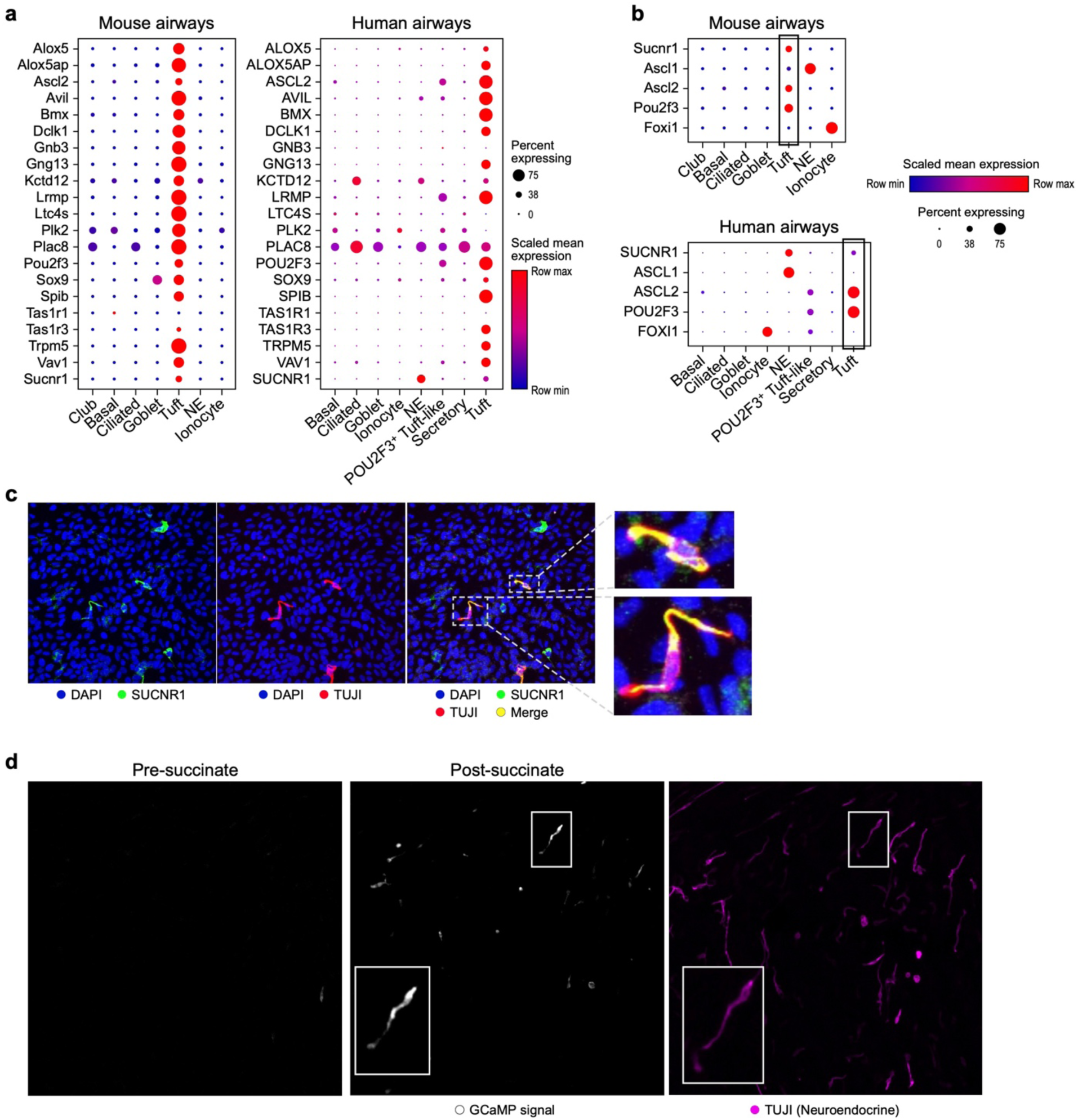
Genes generally associated with murine tuft cell function are distributed across cell types in human bronchial airways. **a,b.** Expression of tuft cell associated genes across different cell types in murine and human airway epithelial cells. Mean expression (dot color,) and percentage of expressing cells (dot size) for murine tuft 1 and tuft 2 cell genes (a, rows) and chemosensory pathway receptors and TFs (b, rows) in either mouse airway (left) or human airway (right) cell subsets. **c.** Chemosensory receptor SUCNR1 is expressed in NE cells in human airways. Antibody staining of SUCNR1 (green) protein in TUJ1^+^ (red) NE cells. Merge - yellow, DAPI- blue. **d**. SUCNR1 expressed in NE cells is responsive to succinate stimulation. TUJ1 antibody staining (NE marker) and GCaMP signal (intracellular calcium wave) in mature ALIs expressing GCaMP6s before and after treatment with succinate. Inset – magnified TUJ1^+^ NE cell displaying intracellular calcium transient.

**Extended Data Figure 6.**
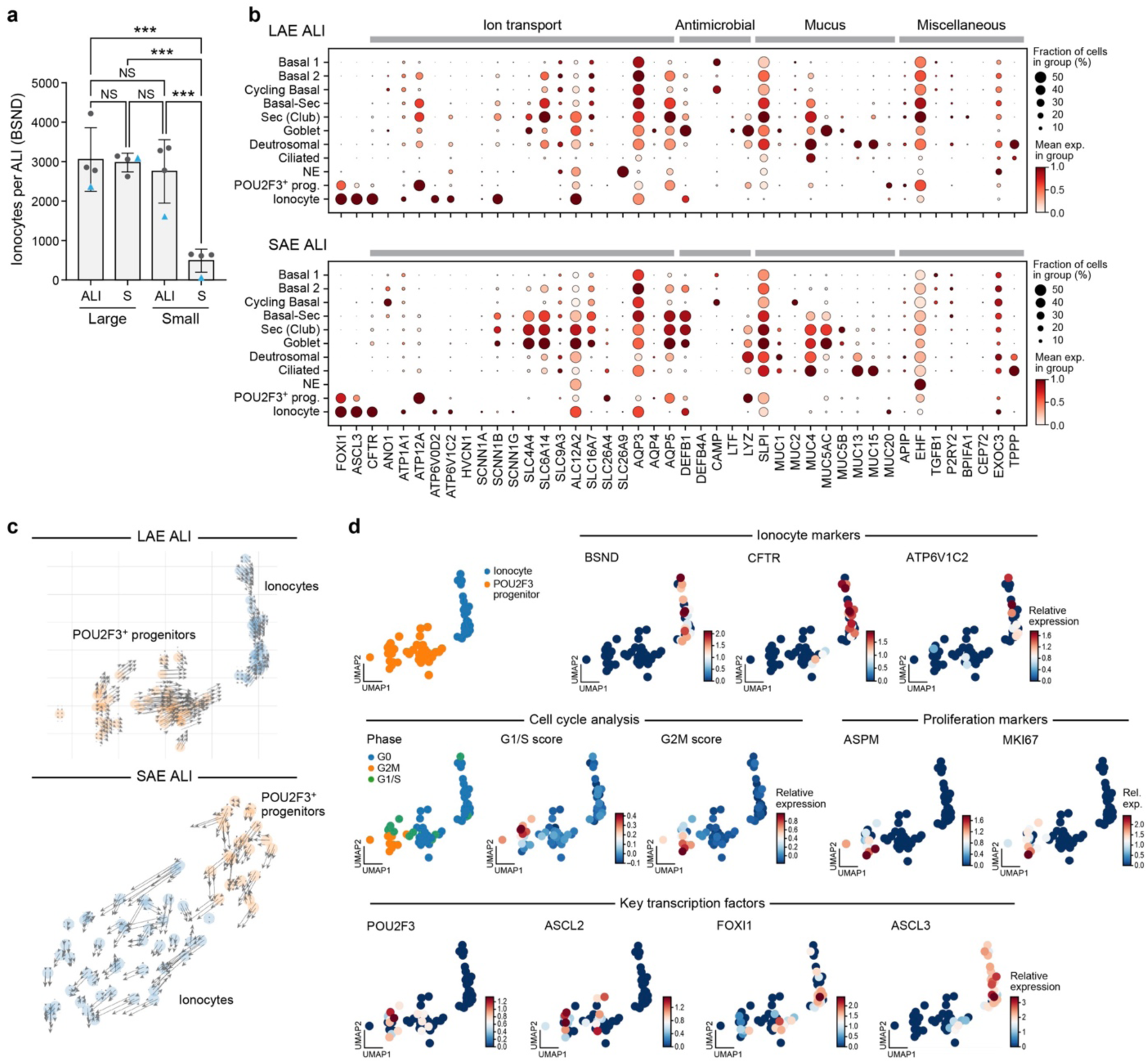
Small airway ALIs grown in appropriate media support the differentiation of ionocytes from POU2F3+ progenitor cells. **a.** Ionocyte differentiation occurs in SAE ALIs under appropriate media conditions. Number of ionocytes (BSND^+^ cells) per ALI (y axis) in large and small airway ALIs grown in regular (‘ALI’) and small airway (‘S’) media (x axis). n=4 (3 biological replicates from 1 human line and 1 replicate from another human line). One way ANOVA. P=0.001 **b**. Expression of CF associated genes in proximal and distal ionocytes in large and small airway ALIs. Mean (dot color, mean expression) and percentage of cells expressing (dot size) genes with CF-associated functions (columns) in each cell subset (rows) from LAE (top) and SAE (bottom) ALIs grown in ALI media. **c.** Differentiation of ionocytes from POU2F3+ progenitors. UMAP embedding of cell profiles (dots) of POU2F3+ progenitors (orange) and ionocytes (blue) from LAE (top) and SAE (bottom) ALI labeled by RNA velocity analysis. **d.** Replicating POU2F3^+^ progenitors and ionocytes in mature LAE ALI cultures . UMAP embedding of cell profiles (dots) of POU2F3+ progenitors (orange) and ionocytes (blue) from LAE ALIs (as in panel c top) colored by cell type annotation (top left), markers of ionocytes (top right), cell cycle phase and phase signature scores (middle left), markers of proliferation (middle right), and the expression of key transcription factor genes (bottom).

**Extended Data Figure 7.**
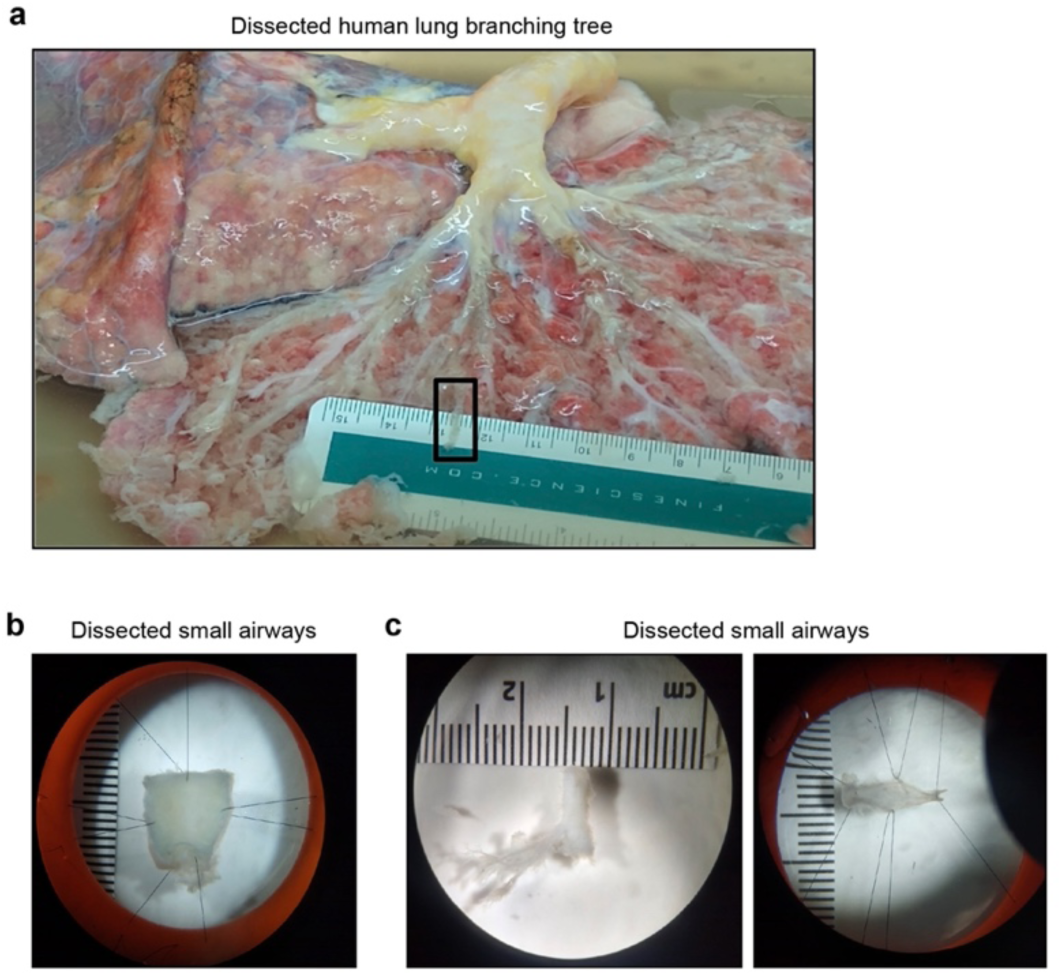
Dissection and mounting of large and small airways for 2-photon imaging. **a.** Dissected human lungs. Black box indicates small airways. **b.** Large airway dissected and mounted for imaging **b.** Two small airways dissected and mounted for imaging.

**Extended Data Figure 8.**
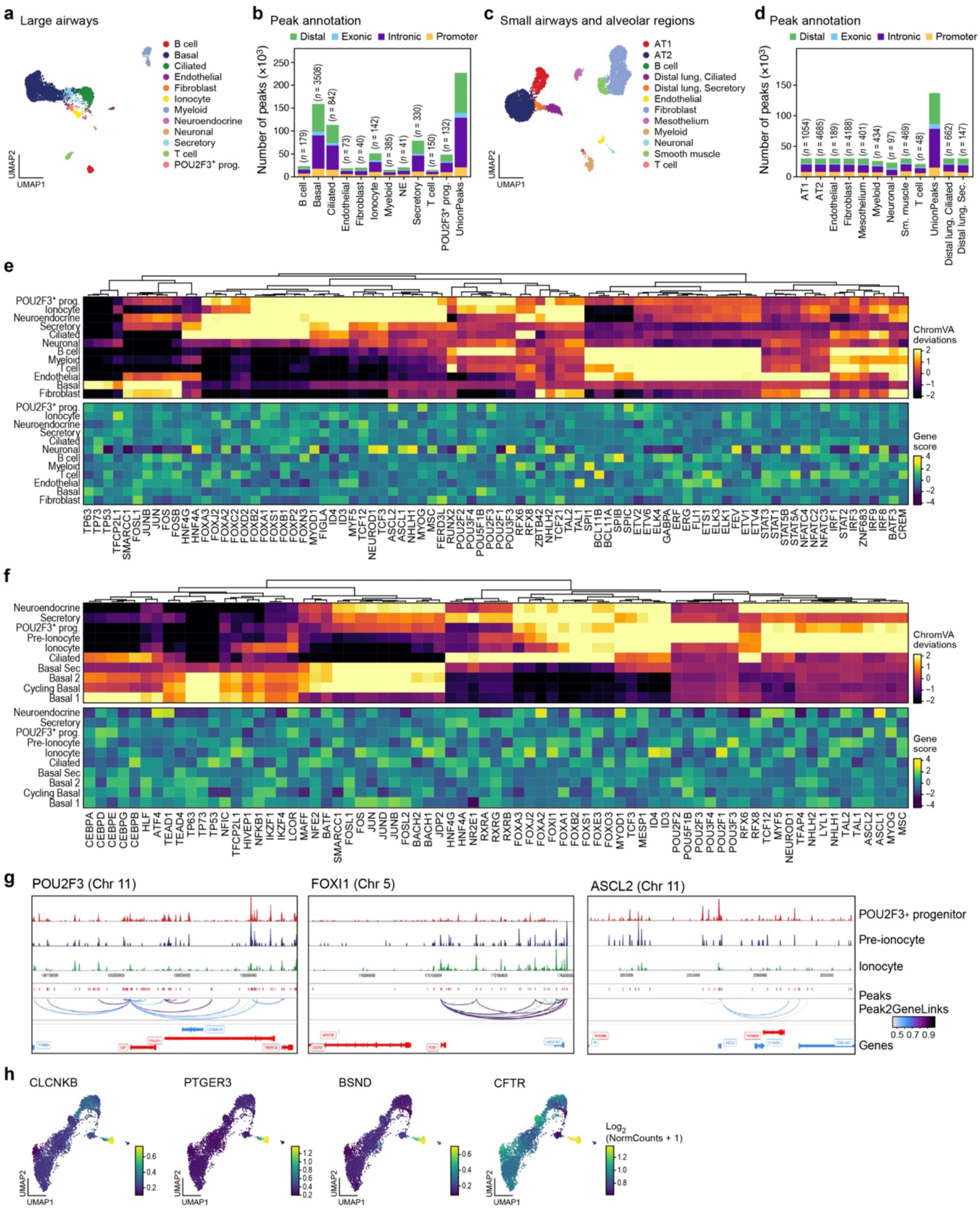
scATACseq of large airways and lung lobe regions. **a-d**. scATAC-seq based cell subsets in the large airways and lung lobes. a,c. UMAP embeddings of scRNA-seq profiles (dots) colored by clustering and annotated *post hoc* for cells from proximal large airways (a) or distal lung lobe regions (c). b,d. Number of peaks (y axis) in each genomic location category (color) for each annotated cell type (x axis) from proximal large airways (b) or distal lung lobe regions (d). **e,f.** Hierarchical clustering of chromVAR top differential TF activity (FDR≤0.01 and, mean differential accessibility ≥0.5) inferred from cognate motif accessibility (top heatmap) and corresponding scaled TF locus accessibility inferred using ArchR gene scores (bottom heatmap) averaged in each cell subset in the large airways (e) and lung lobules (f). **g,h**. Chromatin accessibility at key ionocyte and POU2F3^+^ progenitor loci. g. Aggregate scATAC-seq signal (top three tracks) in POU2F3^+^ progenitors, pre-ionocytes, and ionocyte profiles shown at the *POU2F3* (left), *FOXI1* (middle) and *ASCL2* (right) loci. Bottom three tracks correspond to peaks detected (top), peak to gene links (middle), and gene annotations (bottom). h. UMAP embedding of large airway epithelial scATAC-seq profiles (as in Fig. 2g) colored by ArchR gene score at the gene locus of key ionocyte marker genes.

**Extended Data Figure 9.**
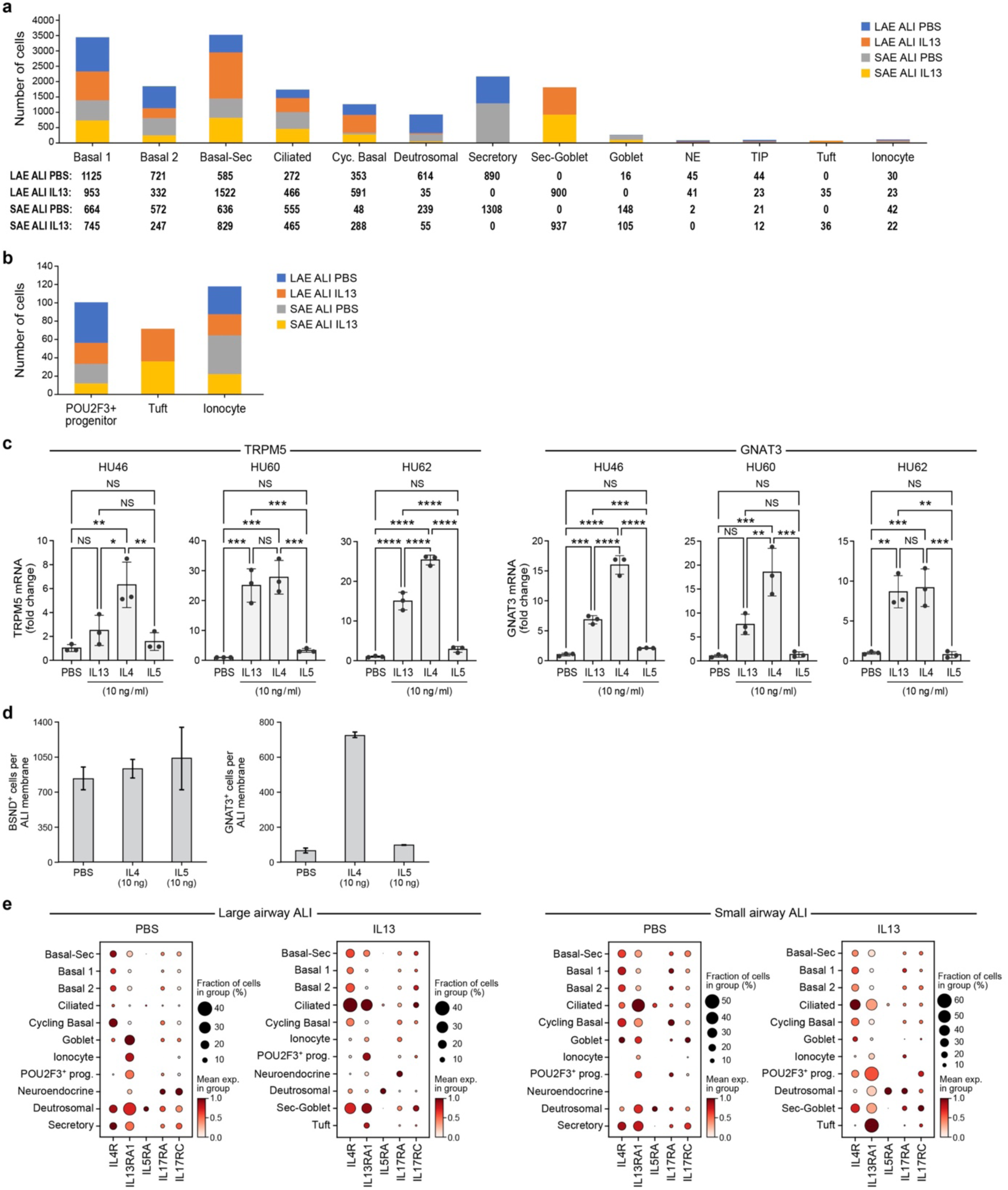
Type 2 signaling promotes tuft cell maturation in ALI culture. **a,b**. IL-13 treatment shifts ALI cell composition. Number of cells (y axis) of each epithelial cell subset (x axis) in LAE and SAE ALI cultures from control (PBS) and IL-13 treated conditions, for all cells (a) and for key rare cell populations (b). **c**. Induction of expression of key tuft cell genes in cytokine treated LAE ALIs. Expression (by qPCR) of mature tuft cell genes in LAE ALIs under control (PBS) conditions or treated with type 2 cytokines IL4, IL5 and IL13 (x axis). Data from ALIs of three human donors (Hu46, Hu60, Hu62) are shown. **d.** Number of antibody-stained cells (y axis) for GNAT3 expressing tuft cells (n=2 (Hu19, Hu67)) and BSND expressing ionocytes (n=3 (Hu19, Hu62, Hu67)) in LAE ALI cultures treated with PBS or IL4 (10ng/ml) or IL5 (10ng/ml) (x axis). **e**. ALI epithelial cells express receptor genes for Type 2 and Type 17 cytokines. Mean normalized expression (dot color) and percentage of expressing cells (dot size) of key cytokine receptor genes (columns) in each epithelial cell subset (rows) in control and IL13-treated LAE and SAE ALIs.

**Extended Data Figure 10.**
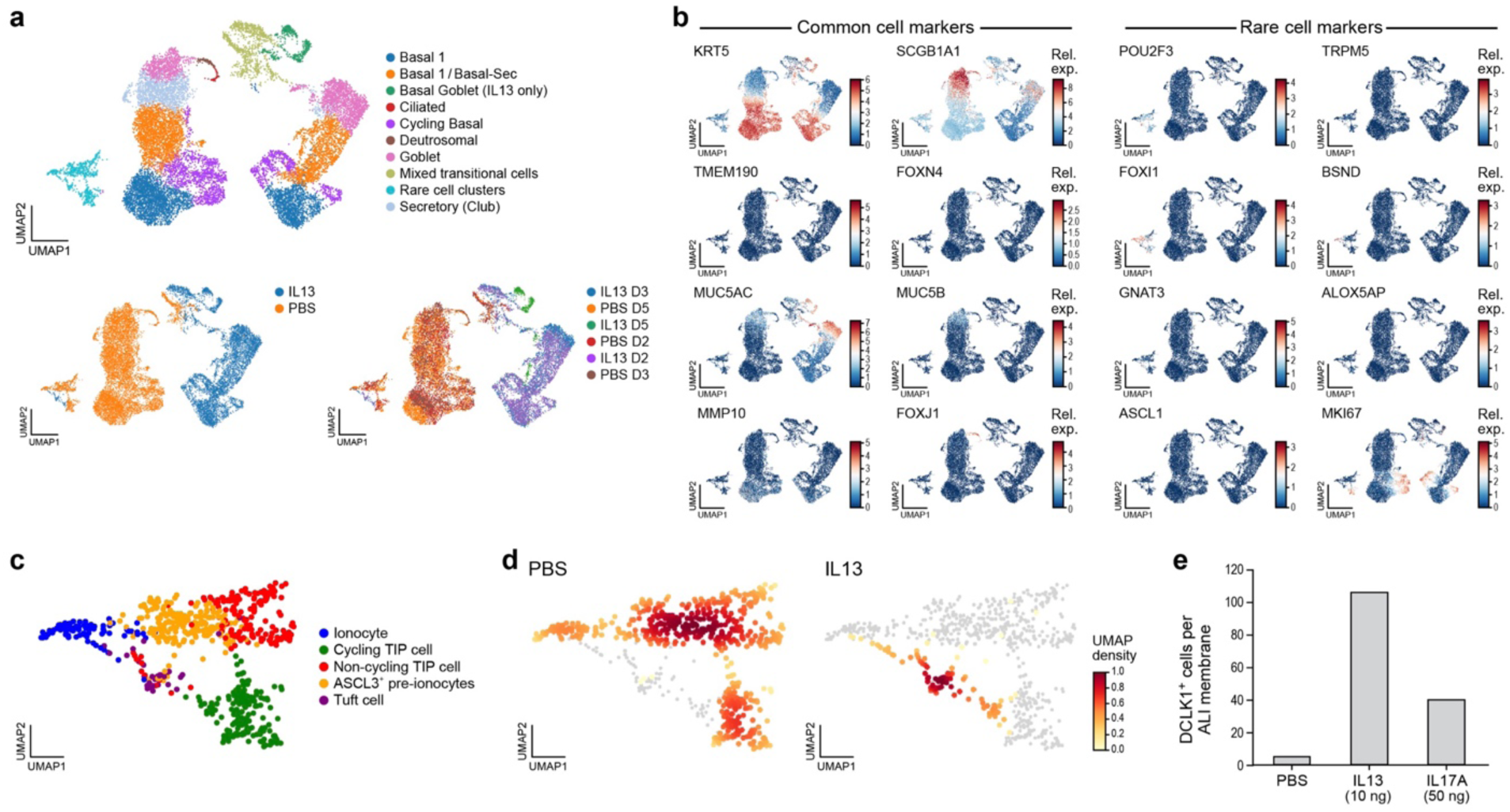
scRNAseq and cell counts of differentiating ALIs following IL13 and IL17A treatment. **a,b.** scRNA-seq of differentiating ALIs. UMAP embedding of cell profiles (dots) from differentiating LAE ALI treated with PBS or IL13 (10ng/µl), colored by cell type annotation (a, top), treatment (a, bottom left), time point (a, bottom right) or expression of markers of common and rare airway epithelial cells types (b). **c,d**. Rare cell differentiation is altered by IL-13 treatment. UMAP embedding of a zoom in of the rare cell type profiles (dots) from both PBS and IL13-treated ALIs labeled by cell type annotation (c) or treatment source (d). **e.** Number of antibody-stained cells (y axis) for DCLK1 expressing tuft cells in SAE ALI treated with PBS or IL13 (10ng/ml) or IL17A (50ng/ml) (x axis) from D3 to D27, n=1.

## ACKNOWLEDGEMENTS

We thank Leslie Gaffney for help with figure preparation, New England Donor Services, the families of donors who consented to the use of donor lungs for research, and the HSCI-CRM Flow Cytometry Core Facility. We thank Asa Segerstolpe, Tommaso Biancalani, Gabriele Scalia and Sanja Vickovic for discussion and experimental input . This work was supported by the Manton Foundation (A.R.) and the Klarman Cell Observatory at the Broad Institute (A.R. and J.R.), Howard Hughes Medical Institute (HHMI), NIH/NHLBI R21HL156124, R56HL157632, R21HL161760, and the Cystic Fibrosis Foundation TSANKO20G0 (A.M.T.), Cystic Fibrosis Foundation 003338L121 (V.S.), Cystic Fibrosis Foundation Lin19F0 (B.L.), NIH 5U24HL148865-04/OS00000379 (V.S.), a LungMAP2 Pilot Grant under NIH 1U24HL148865-01(A.W.), the Chan Zuckerberg Human Cell Atlas Initiative (J.R.), and Mike Toth Head and Neck cancer funds (SVS). A.R. was an Investigator of the Howard Hughes Medical Institute. This publication is part of the Human Cell Atlas – www.humancellatlas.org/publications/.

## REFERENCES

1. Goldfarbmuren, K. C. et al. Dissecting the cellular specificity of smoking effects and reconstructing lineages in the human airway epithelium. Nat. Commun. 11, 2485 (2020).

2. Adams, T. S. et al. Single-cell RNA-seq reveals ectopic and aberrant lung-resident cell populations in idiopathic pulmonary fibrosis. Science Advances vol. 6 eaba1983 Preprint at 10.1126/sciadv.aba1983 (2020).

3. Braga, F. A. V. et al. A cellular census of human lungs identifies novel cell states in health and in asthma. Nat. Med.

4. Travaglini, K. J. et al. A molecular cell atlas of the human lung from single-cell RNA sequencing. Nature 587, 619–625 (2020).

5. Deprez, M. et al. A Single-Cell Atlas of the Human Healthy Airways. Am. J. Respir. Crit. Care Med. 202, 1636–1645 (2020).

6. Plasschaert, L. W. et al. A single-cell atlas of the airway epithelium reveals the CFTR-rich pulmonary ionocyte. Nature 560, 377–381 (2018).

7. Montoro, D. T. et al. A revised airway epithelial hierarchy includes CFTR-expressing ionocytes. Nature (2018) doi:10.1038/s41586-018-0393-7.

8. Madissoon, E. et al. scRNA-seq assessment of the human lung, spleen, and esophagus tissue stability after cold preservation. Genome Biol. 21, 1 (2019).

9. Carraro, G. et al. Single-Cell Reconstruction of Human Basal Cell Diversity in Normal and Idiopathic Pulmonary Fibrosis Lungs. Am. J. Respir. Crit. Care Med. 202, 1540–1550 (2020).

10. Ruiz García, S., et al. Novel dynamics of human mucociliary differentiation revealed by single-cell RNA sequencing of nasal epithelial cultures. Development 146, (2019).

11. Wang, A. et al. Single-cell multiomic profiling of human lungs reveals cell-type-specific and age-dynamic control of SARS-CoV2 host genes. Elife 9, (2020).

12. Yuan, F. et al. Transgenic ferret models define pulmonary ionocyte diversity and function. Nature 621, 857–867 (2023).

13. Cai, Q. et al. Sonic Hedgehog Signaling is Essential for Pulmonary Ionocyte Specification in Human and Ferret Airway Epithelia. Am. J. Respir. Cell Mol. Biol. (2023) doi:10.1165/rcmb.2022-0280OC.

14. Lei, L. et al. CFTR-rich ionocytes mediate chloride absorption across airway epithelia. J. Clin. Invest. 133, (2023).

15. Žilionis, R., et al. A single-cell atlas of the airway epithelium reveals the CFTR-rich pulmonary ionocyte. Nature (2018).

16. Sikkema, L. et al. An integrated cell atlas of the lung in health and disease. Nat. Med. 29, 1563–1577 (2023).

17. Dimaio, M. F., Dische, R., Gordon, R. E. & Kattan, M. Alveolar brush cells in an infant with desquamative interstitial pneumonitis. Pediatr. Pulmonol. 4, 185–191 (1988).

18. Gordon, R. E. y., E. y Gordon, R. & Kattan, M. Absence of Cilia and Basal Bodies with Predominance of Brush Cells in the Respiratory Mucosa from a Patient with Immotile Cilia Syndrome. Ultrastructural Pathology vol. 6 45–49 Preprint at 10.3109/01913128409016664 (1984).

19. Melms, J. C. et al. A molecular single-cell lung atlas of lethal COVID-19. Nature 595, 114– 119 (2021).

20. Gerbe, F. et al. Intestinal epithelial tuft cells initiate type 2 mucosal immunity to helminth parasites. Nature 529, 226–230 (2016).

21. Schneider, C. et al. A Metabolite-Triggered Tuft Cell-ILC2 Circuit Drives Small Intestinal Remodeling. Cell 174, 271–284.e14 (2018).

22. Howitt, M. R. et al. Tuft cells, taste-chemosensory cells, orchestrate parasite type 2 immunity in the gut. Science 351, 1329–1333 (2016).

23. von Moltke, J., Ji, M., Liang, H.-E. & Locksley, R. M. Tuft-cell-derived IL-25 regulates an intestinal ILC2–epithelial response circuit. Nature 529, 221–225 (2015).

24. Okuda, K. et al. Secretory Cells Dominate Airway CFTR Expression and Function in Human Airway Superficial Epithelia. Am. J. Respir. Crit. Care Med. 203, 1275–1289 (2021).

25. Yamashita, J., Ohmoto, M., Yamaguchi, T., Matsumoto, I. & Hirota, J. Skn-1a/Pou2f3 functions as a master regulator to generate Trpm5-expressing chemosensory cells in mice. PLoS One 12, e0189340 (2017).

26. Basil, M. C. et al. Human distal airways contain a multipotent secretory cell that can regenerate alveoli. Nature 604, 120–126 (2022).

27. Kadur Lakshminarasimha Murthy, P., et al. Human distal lung maps and lineage hierarchies reveal a bipotent progenitor. Nature 604, 111–119 (2022).

28. Sikkema, L. et al. An integrated cell atlas of the human lung in health and disease. bioRxiv 2022.03.10.483747 (2022) doi:10.1101/2022.03.10.483747.

29. Shah, A. S. Motile cilia of human airway are chemosensory. Science 325, 1131–1134 (2009).

30. Gu, X. et al. Chemosensory Functions for Pulmonary Neuroendocrine Cells. Am. J. Respir. Cell Mol. Biol. 50, 637–646 (2014).

31. Yu, W., Moninger, T. O., Rector, M. V., Stoltz, D. A. & Welsh, M. J. Pulmonary neuroendocrine cells sense succinate to stimulate myoepithelial cell contraction. Dev. Cell 57, 2221–2236.e5 (2022).

32. Wang, G. et al. Characterization of an immortalized human small airway basal stem/progenitor cell line with airway region-specific differentiation capacity. Respir. Res. 20, 196 (2019).

33. Bergen, V., Lange, M., Peidli, S., Wolf, F. A. & Theis, F. J. Generalizing RNA velocity to transient cell states through dynamical modeling. Nat. Biotechnol. 38, 1408–1414 (2020).

34. Schep, A. N., Wu, B., Buenrostro, J. D. & Greenleaf, W. J. chromVAR: inferring transcription-factor-associated accessibility from single-cell epigenomic data. Nat. Methods 14, 975–978 (2017).

35. Ma, S. et al. Chromatin Potential Identified by Shared Single-Cell Profiling of RNA and Chromatin. Cell 183, 1103–1116.e20 (2020).

36. Suzukawa, M. et al. Epithelial cell-derived IL-25, but not Th17 cell-derived IL-17 or IL-17F, is crucial for murine asthma. J. Immunol. 189, 3641–3652 (2012).

37. Ualiyeva, S., et al. Tuft cell-produced cysteinyl leukotrienes and IL-25 synergistically initiate lung type 2 inflammation. *Sci Immunol* 6, eabj0474 (2021).

38. McGinty, J. W. et al. Tuft-Cell-Derived Leukotrienes Drive Rapid Anti-helminth Immunity in the Small Intestine but Are Dispensable for Anti-protist Immunity. Immunity 52, 528– 541.e7 (2020).

39. Kotas, M. E., et al. IL-13-programmed airway tuft cells produce PGE2, which promotes CFTR-dependent mucociliary function. *JCI Insight* 7, (2022).

40. Kohanski, M. A. et al. Solitary chemosensory cells are a primary epithelial source of IL-25 in patients with chronic rhinosinusitis with nasal polyps. J. Allergy Clin. Immunol. 142, 460–469.e7 (2018).

41. Beale, J. et al. Rhinovirus-induced IL-25 in asthma exacerbation drives type 2 immunity and allergic pulmonary inflammation. Sci. Transl. Med. 6, 256ra134 (2014).

42. Tojima, I. et al. Evidence for the induction of Th2 inflammation by group 2 innate lymphoid cells in response to prostaglandin D2 and cysteinyl leukotrienes in allergic rhinitis. Allergy 74, 2417–2426 (2019).

43. Lee, R. J. et al. Bacterial d-amino acids suppress sinonasal innate immunity through sweet taste receptors in solitary chemosensory cells. Sci. Signal. 10, (2017).

44. Sell, E. A., Ortiz-Carpena, J. F., Herbert, D. R. & Cohen, N. A. Tuft cells in the pathogenesis of chronic rhinosinusitis with nasal polyps and asthma. Ann. Allergy Asthma Immunol. 126, 143–151 (2021).

45. Lee, R. J. et al. Bitter and sweet taste receptors regulate human upper respiratory innate immunity. J. Clin. Invest. 124, 1393–1405 (2014).

46. Alladina, J., et al. A human model of asthma exacerbation reveals transcriptional programs and cell circuits specific to allergic asthma. *Sci Immunol* 8, eabq6352 (2023).

47. Jackson, N. D. et al. Single-Cell and Population Transcriptomics Reveal Pan-epithelial Remodeling in Type 2-High Asthma. Cell Rep. 32, 107872 (2020).

48. Huang, Y.-H. et al. POU2F3 is a master regulator of a tuft cell-like variant of small cell lung cancer. 915–928 (2018).

49. Rane, C. K. et al. Development of solitary chemosensory cells in the distal lung after severe influenza injury. Am. J. Physiol. Lung Cell. Mol. Physiol. 316, L1141–L1149 (2019).

50. Barr, J. et al. Injury-induced pulmonary tuft cells are heterogenous, arise independent of key Type 2 cytokines, and are dispensable for dysplastic repair. Elife 11, (2022).

51. Zhang, Y. et al. Immune Cell Production of Interleukin 17 Induces Stem Cell Features of Pancreatic Intraepithelial Neoplasia Cells. Gastroenterology 155, 210–223.e3 (2018).

52. Hackett, T.-L. et al. Intrinsic phenotypic differences of asthmatic epithelium and its inflammatory responses to respiratory syncytial virus and air pollution. Am. J. Respir. Cell Mol. Biol. 45, 1090–1100 (2011).

53. Xiao, C. et al. Defective epithelial barrier function in asthma. J. Allergy Clin. Immunol. 128, 549–56.e1–12 (2011).

54. Gras, D. et al. An ex vivo model of severe asthma using reconstituted human bronchial epithelium. J. Allergy Clin. Immunol. 129, 1259–1266.e1 (2012).

55. Ordovas-Montanes, J. et al. Allergic inflammatory memory in human respiratory epithelial progenitor cells. Nature 560, 649–654 (2018).

56. Shivaraju, M. et al. Airway stem cells sense hypoxia and differentiate into protective solitary neuroendocrine cells. Science 371, 52–57 (2021).

57. Conchola, A. S. et al. Regionally distinct progenitor cells in the lower airway give rise to neuroendocrine and multiciliated cells in the developing human lung. Proc. Natl. Acad. Sci. U. S. A. 120, e2210113120 (2023).

58. Feldman, M. B., Wood, M., Lapey, A. & Mou, H. SMAD Signaling Restricts Mucous Cell Differentiation in Human Airway Epithelium. Am. J. Respir. Cell Mol. Biol. 61, 322–331 (2019).

59. Stuart, T. et al. Comprehensive Integration of Single-Cell Data. Cell 177, 1888–1902.e21 (2019).

60. Tirosh, I. et al. Dissecting the multicellular ecosystem of metastatic melanoma by single-cell RNA-seq. Science 352, 189–196 (2016).

61. La Manno, G. et al. RNA velocity of single cells. Nature 560, 494–498 (2018).

